# YgfB increases β-lactam resistance in *Pseudomonas aeruginosa* by counteracting AlpA-mediated *ampDh3* expression

**DOI:** 10.1101/2022.07.05.498809

**Authors:** Ole Eggers, Fabian Renschler, Lydia Anita Michalek, Noelle Wackler, Elias Walter, Fabian Smollich, Kristina Klein, Michael Sonnabend, Valentin Egle, Angel Angelov, Christina Engesser, Marina Borisova, Christoph Mayer, Monika Schütz, Erwin Bohn

## Abstract

YgfB-mediated β-lactam resistance was recently identified in multi drug resistant *Pseudomonas aeruginosa. We* show that YgfB upregulates expression of the β-lactamase AmpC by repressing the function of the regulator of the programmed cell death pathway AlpA. In response to DNA damage, the antiterminator AlpA induces expression of the *alpBCDE* autolysis genes and of the peptidoglycan amidase AmpDh3. YgfB interacts with AlpA and represses the *ampDh3* expression.

Thus, YgfB indirectly prevents AmpDh3 from reducing the levels of cell wall-derived 1,6-anhydro-N-acetylmuramyl-peptides, required to induce the transcriptional activator AmpR in promoting the *ampC* expression and β-lactam resistance. Ciprofloxacin-mediated DNA damage induces AlpA-dependent production of AmpDh3 as previously shown, which should reduce β-lactam resistance. YgfB, however, counteracts the β-lactam enhancing activity of ciprofloxacin by repressing *ampDh3* expression and lowering the benefits of this drug combination.

Altogether, YgfB represents a new player in the complex regulatory network of AmpC regulation.

## Introduction

The rise in multi drug resistance (MDR) represents a major public health issue due to declining options for the treatment of bacterial infections. This poses a large problem particularly for nosocomial infections with *Pseudomonas aeruginosa*, which is one of the most important Gram-negative pathogens and a major cause of pneumonia, urinary tract infections, wound infections and blood stream infections. MDR in *P. aeruginosa* is constituted by various intrinsic and acquired antibiotic resistance mechanisms. High intrinsic resistance is mainly the result of a very low permeability of the outer membrane^1^ and the inducible expression of efflux pumps^2^. High-level resistance to β-lactams, including resistance to carbapenems and cephalosporins, is mediated by an upregulation of the chromosomally-encoded cephalosporinase AmpC^2^. While AmpC-levels are low in wild type *P. aeruginosa* strains, they are often strongly increased in clinical isolates. Derepression of *ampC* frequently occurs due to loss of function mutations in *ampR, ampD, or dacB, which* encode the transcriptional regulator AmpR, a cytoplasmic muropeptide amidase^3, 4^, and penicillin-binding protein 4 (PBP4), respectively^5^.

Additionally, *ampC* expression can be induced by specific β-lactam antibiotics and β-lactamase inhibitors. This results in *P. aeruginosa* strains that are resistant to most β-lactam antibiotics^6^. AmpR regulates *ampC* expression by sensing the relative amounts of peptidoglycan (PG) recycling metabolites and synthesis precursors^7^. When binding the final soluble PG synthesis precursor UDP-N-acetylmuramic acid pentapeptide (UDP-MurNAc-5P), AmpR represses a subset of genes including *ampC*^8^. On the contrary, 1,6-anhydro-N-acetyl-MurNAc-peptides (anhMurNAc-peptides) originating from the turnover and recycling of the PG cell wall^9^, can displace UDP-MurNAc-5P from AmpR. This allows AmpR to transactivate various promoters such as the *ampC* promoter, causing high levels of AmpC-β-lactamase. Interestingly, those mutations in *ampD* or *dacB* that cause derepression of *ampC* give rise to high intracellular pool levels of anhMurNAc-peptides such as 1,6-anhMurNAc-L-alanyl-D-glutamyl-meso-diaminopimelic acid-D-alanyl-D-alanine, i.e., anhMurNAc-pentapeptide (anhMurNAc-5P)^7^.

The MDR *P. aeruginosa* strain ID40 is a bloodstream isolate carrying a loss of function mutation in *dacB*, hence exhibits high intrinsic β-lactamase activity^10, 11^. In a previous study we used an ID40 transposon (Tn) library to identify genes involved in β-lactam resistance. For this purpose, a Tn library was grown in the presence of cefepime, meropenem or without antibiotics and subsequently transposon directed insertion sequencing (TraDIS) was performed^10^. Among the identified genes we found the so far uncharacterized gene *ygfB*, whose expression significantly increases the levels of AmpC, β-lactamase activity and consequently, resistance to various classes of β-lactam antibiotics such as carbapenems, monobactams, and 3^rd^ and 4^th^ generation cephalosporins^10^. YgfB belongs to the UPF0149 family of proteins with orthologs found in many γ-proteobacteria **(Fig. S1)**. YgfB consists of seven α-helices and the structure analysis of the *Haemophilus influenzae* ortholog suggests that YgfB forms a homodimer (PDB ID 1IZM^12^). To this point and to our knowledge, nothing else is currently known about the function of YgfB homologous proteins.

In the present study, we unraveled how YgfB contributes to β-lactam resistance. We show that in *P. aeruginosa* YgfB decreases the production levels of the amidase AmpDh3 by repressing the activity of the antiterminator protein AlpA, which controls the expression of AmpDh3. This role of YgfB seems to be conserved across different clinical isolates of *P. aeruginosa*. Furthermore, we provide a molecular explanation to why the ciprofloxacin/β-lactam combination therapy is ineffective in many cases of *P. aeruginosa* infections.

## Results

### Transcriptome analysis reveals specific effects of YgfB on gene expression

To investigate the role of YgfB in antibiotic resistance, a transcriptome analysis was performed using next generation sequencing with mRNAs isolated from ID40 and the isogenic *ygfB* null mutant ID40Δ*ygfB*. As depicted in **Table 1**, the mRNA expression of only few genes was significantly altered following *ygfB* deletion, indicating a very specific effect of YgfB on the transcriptional level.

**Table 1:**
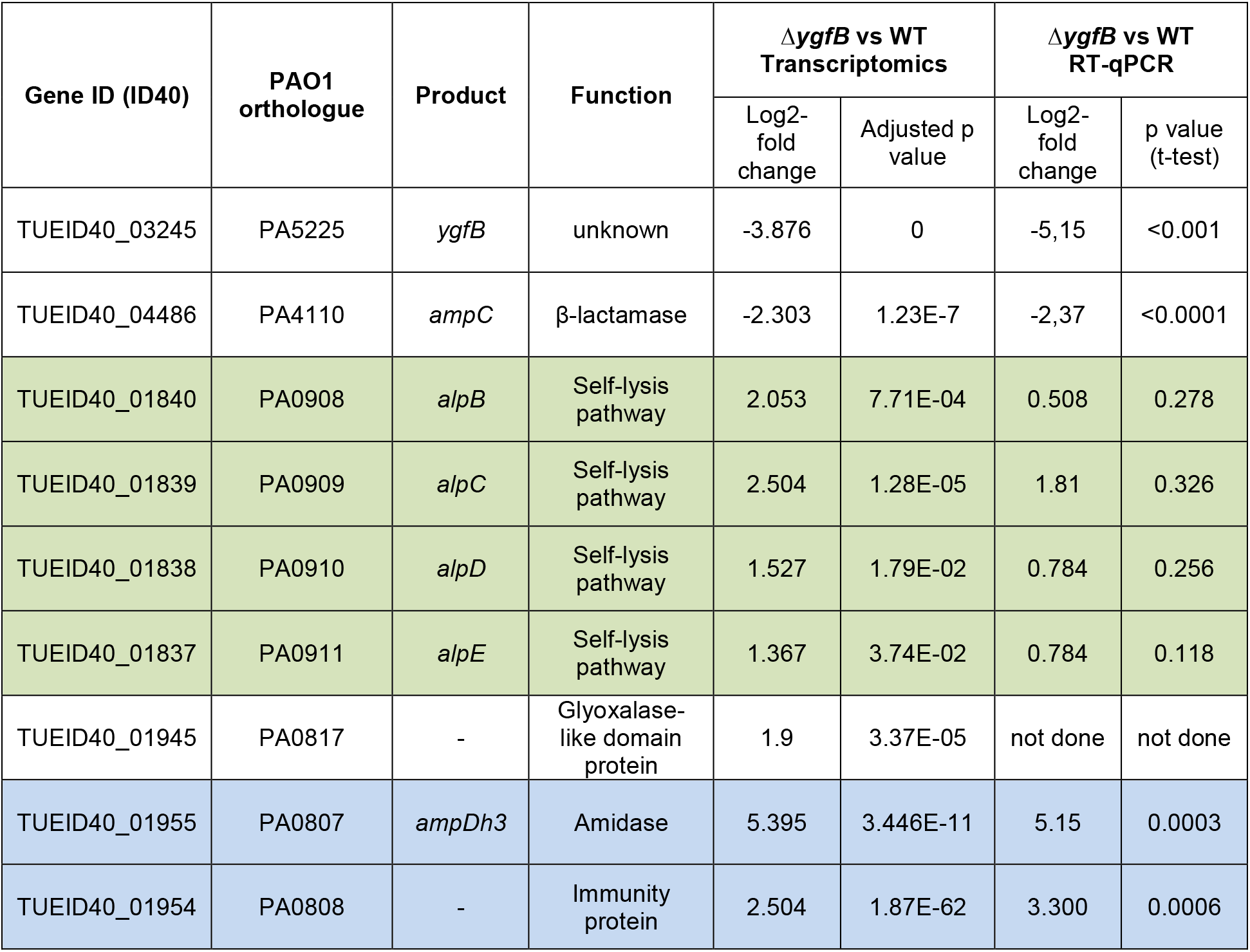
Transcriptomic analyses and validation by RT-qPCR. Complete list of genes with significant changes in mRNA expression due to *ygfB* deletion. Transcriptomics *n*=3 with data analysis as described in methods section, qRT-PCR *n*=3-6, two-tailed unpaired t-test.

| Gene ID (ID40) | PAO1 orthologue | Product | Function | $\Delta ygfB$ vs WT Transcriptomics | | $\Delta ygfB$ vs WT RT-qPCR | |
| --- | --- | --- | --- | --- | --- | --- | --- |
|  |  |  |  | Log2-fold change | Adjusted p value | Log2-fold change | p value (t-test) |
| TUEID40_03245 | PA5225 | *ygfB* | unknown | -3.876 | 0 | -5,15 | <0.001 |
| TUEID40_04486 | PA4110 | *ampC* | β-lactamase | -2.303 | 1.23E-7 | -2,37 | <0.0001 |
| TUEID40_01840 | PA0908 | *alpB* | Self-lysis pathway | 2.053 | 7.71E-04 | 0.508 | 0.278 |
| TUEID40_01839 | PA0909 | *alpC* | Self-lysis pathway | 2.504 | 1.28E-05 | 1.81 | 0.326 |
| TUEID40_01838 | PA0910 | *alpD* | Self-lysis pathway | 1.527 | 1.79E-02 | 0.784 | 0.256 |
| TUEID40_01837 | PA0911 | *alpE* | Self-lysis pathway | 1.367 | 3.74E-02 | 0.784 | 0.118 |
| TUEID40_01945 | PA0817 | - | Glyoxalase-like domain protein | 1.9 | 3.37E-05 | not done | not done |
| TUEID40_01955 | PA0807 | *ampDh3* | Amidase | 5.395 | 3.446E-11 | 5.15 | 0.0003 |
| TUEID40_01954 | PA0808 | - | Immunity protein | 2.504 | 1.87E-62 | 3.300 | 0.0006 |

Deletion of *ygfB* resulted in the downregulation of only one gene, *ampC*, encoding the β-lactamase, and the upregulation of seven other genes including the *ampDh3*-TUEID40_01954 operon, a gene encoding a glyoxalase-like protein and the *alpBCDE* genes. Together with its paralogues AmpD and AmpDh2, AmpDh3 belongs to the family of amidases that are able to remove the peptide stem from peptidoglycan intermediates^13^. All three amidases were described to contribute to β-lactam resistance^14^. AlpA was recently described to be an antiterminator that transcriptionally regulates the *alpBCDE* cluster, encoding a self-lysis pathway, and the *ampDh3* operon. The activation of the self-lysis pathway can occur in a subset of cells in response to DNA damage, is lethal to the individual cells in which it occurs and might be required to enhance pathogenicity during lung infection^15^. Since it was shown that AlpA specifically upregulates the expression of both the *alpBCDE* cluster and *ampDh3*, we speculated that YgfB could be a negative regulator of AlpA-mediated transcription^15, 16, 17^. Based on the amidase function of AmpDh3 we hypothesized that *ygfB* deletion could increase the AmpDh3 levels, thereby leading to a reduction in the levels of anhMurNAc-peptides. This would in turn lead to a reduction in the transcriptional activation of *ampC* by AmpR and an increased sensitivity to β-lactam antibiotics.

### Validation of transcriptomics results by RT-qPCR and Western blots

The transcriptomics results were validated by RT-qPCR and Western blot analysis. For this purpose, we compared mRNA and protein levels between the strains ID40, ID40Δ*ygfB* and ID40Δ*ygfB* complemented with an genomic copy of *ygfB* under the control of a rhamnose inducible promoter *(ΔygfB::rha-ygfB)* **(Fig. 1a and b)**. We found that *ygfB* deletion significantly reduced the expression of *ampC* and increased the expression of the operon encoding for *ampDh3* and TUEID40_01954 **(Fig. 1a, Table 1)**. Complementation was achieved by the addition of various concentrations of rhamnose, leading to increased *ygfB* and *ampC* expression and a parallel decrease in *ampDh3* and TUEID40_01954 expression. The expression of the *ampDh3* paralogues *ampD* and *ampDh2* was not affected by the *ygfB* deletion. While we could see a clear effect of *ygfB* deletion on the mRNA expression of the *alpBCDE* genes in transcriptomic analysis, only a subtle, non-significant impact of YgfB expression on the *alpBCDE* cluster could be observed for the strain ID40 when trying to validate the results by RT-qPCR. **(Fig. S2, Table 1)**.

**Fig. 1:**
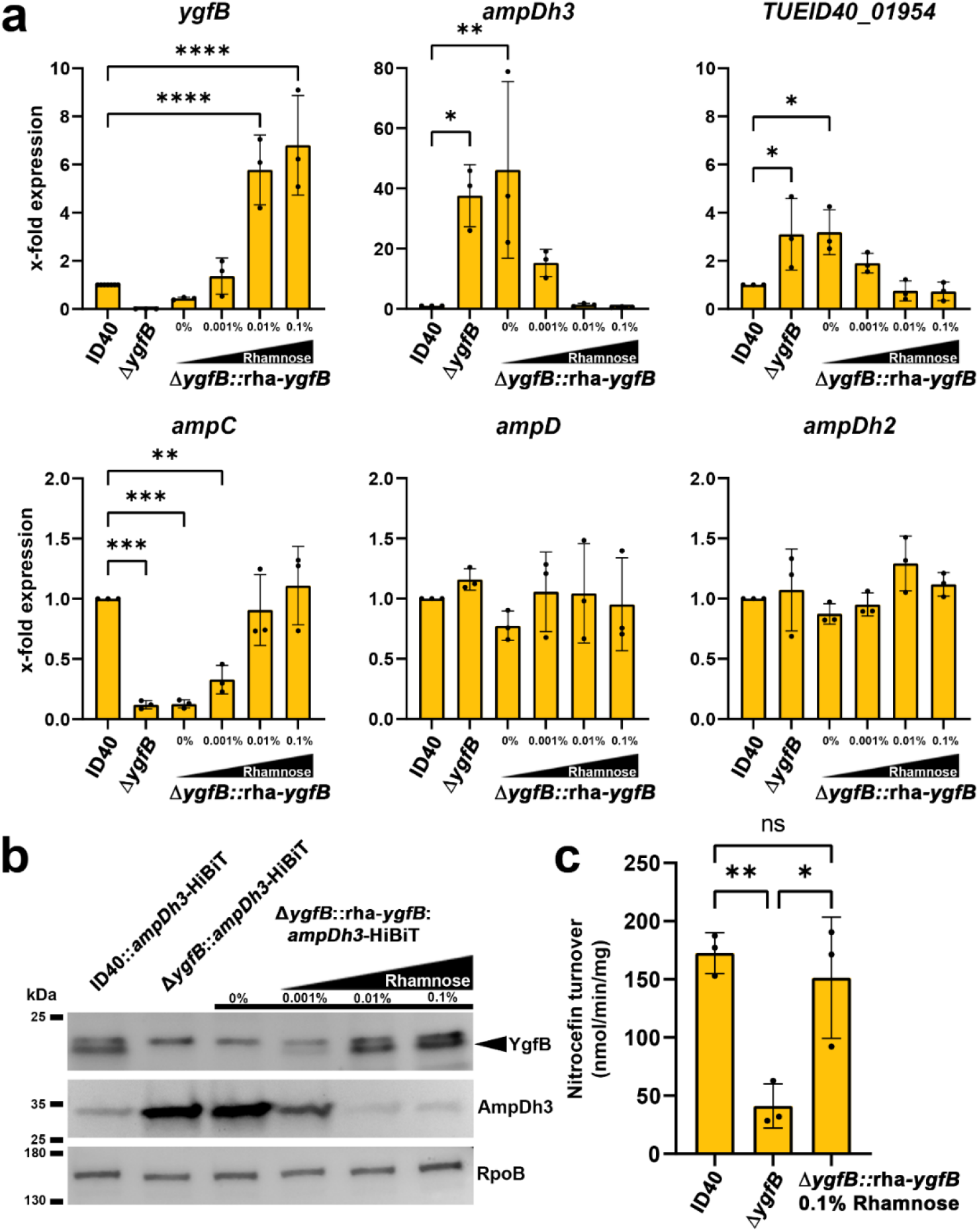
Validation of transcriptomic data. (a) RNA was isolated from the indicated strains and RT-qPCR performed. Data depict the mean and SD of the x-fold difference in mRNA expression compared to ID40 of *n*=3 individual experiments. Asterisks depict significant differences (*p<0.05, **p<0.01, ****p<0.001; one-way ANOVA, Dunnett’s multiple comparison comparing to ID40). (b) Whole cell lysates of the indicated strains were used for SDS-PAGE and Western blots. As primary antibodies, anti-YgfB or as a loading control anti-RpoB antibodies and as secondary antibody anti-IgG-HRP antibodies were used and detection was done using ECL. For determination of AmpDh3, recombinant LgBiT was used. LgBiT binds to HiBiT resulting in a functional luciferase. The cleavage of the substrate furimazine leads to detectable chemiluminescence. Data are representative of three independent experiments (c) Whole cell lysates from the indicated strains were used to determine β-lactamase activity using a nitrocefin assay. Data depict the mean and SD of nitrocefin turnover for *n*=3 individual experiments. Asterisks depict significant differences (ns p>0.05, *p<0.05, **p<0.01, ****p<0.0001; one-way ANOVA, Tukey’s multiple comparisons).

To investigate the modulation of AmpDh3 also at the protein level, the *ampDh3* gene was replaced via allelic exchange by a gene encoding AmpDh3 fused to the HiBiT fragment of the split luciferase protein^18^. This yielded the strains ID40::*ampDh3*-HiBiT, *ID40ΔygfB::ampDh3-* HiBiT, and ID40Δ*ygfB*::rha-*ygfB*::*ampDh3*-HiBiT. Western blot analysis revealed that the deletion of *ygfB* increased the protein levels of AmpDh3. Complementation experiments confirmed that YgfB levels were negatively associated with that of AmpDh3 **(Fig. 1b**). It should be noted that the antibodies we used for detection of YgfB produced an unspecific band very close to YgfB but at a slightly higher molecular weight. Semiquantification of AmpDh3-HiBiT production by using a luciferase assay (**Fig S3a, right panel**) indicates an increase of more than 30-fold upon *ygfB* deletion compared to ID40. Reintroduction and expression of a rhamnose inducible copy of *ygfB* in the null mutant strain reduced AmpDh3 production by 98% (**Fig. S3a**). Semiquantification of YgfB and AmpDh3 based on the western blots in Figure 1 is also provided in **Fig. S3a**. By employing a β-lactamase assay with the chromogenic substrate nitrocefin, we could furthermore show that the reduction of *ampC* expression caused by the deletion of *ygfB* actually translated in a reduced β-lactamase activity. This effect could be reversed by the induction of *ygfB* expression via the addition of rhamnose in the strain Δ*ygfB*::rha-*ygfB* **(Fig. 1c).**

### YgfB upregulates AmpC by repressing *ampDh3* expression

Our previous experiments have shown that *ygfB* deletion has opposite effects on *ampC* and *ampDh3* expression. Therefore, we asked whether the increased expression levels of AmpDh3 could be causative of the reduction in *ampC* expression and consequently the AmpC protein levels in ID40Δ*ygfB*. To challenge this hypothesis, the expression of *ygfB* was induced in the strain ID40Δ*ygfB*::rha-*ygfB* by the addition of 0.1% rhamnose and mRNA transcripts of *ygfB, ampC* and *ampDh3* were quantified by RT-qPCR at different time points **(Fig. 2a)**. Induction of *ygfB* led to a fast decrease of *ampDh3* expression and a time-delayed increase in *ampC* mRNA expression. To confirm these findings, the strains ID40::*ampDh*3-HiBiT, ID40Δ*ygfB*::*ampDh3*-HiBiT and ID40Δ*ygfB*::rha-*ygfB*::*ampDh3-HiBiT* were used for Western blot analysis. A negative association between YgfB and AmpDh3 protein levels could also be observed **(Fig. 2b)**. Semiquantification of the YgfB and AmpDh3-HiBiT bands of the Western blots as well as additional semiquantification of AmpDh3-HiBiT levels in cell lysates using a luciferase assay are shown in **Fig. S3b**. Taken together, these data further supported the hypothesis that YgfB might act as a repressor of *ampDh3* expression, and that the levels of AmpDh3 might influence β-lactam resistance in ID40. To challenge these assumptions further, *ampDh3* and *ygfB*/*ampDh3* deletion mutants were generated and RT-qPCR was performed to analyze *ygfB*, *ampDh3* and *ampC* expression levels. As shown in **Fig. 2c**, the concurrent deletion of *ygfB* and *ampDh3* restored *ampC* expression, as well as β-lactamase activity (**Fig. 2d**). In contrast, *ampDh3* deletion had no impact on the expression of TUEID40_01954. This demonstrates that the repression of *ampDh3* expression by YgfB is required to achieve high *ampC* expression in the *P. aeruginosa* strain ID40.

**Fig. 2:**
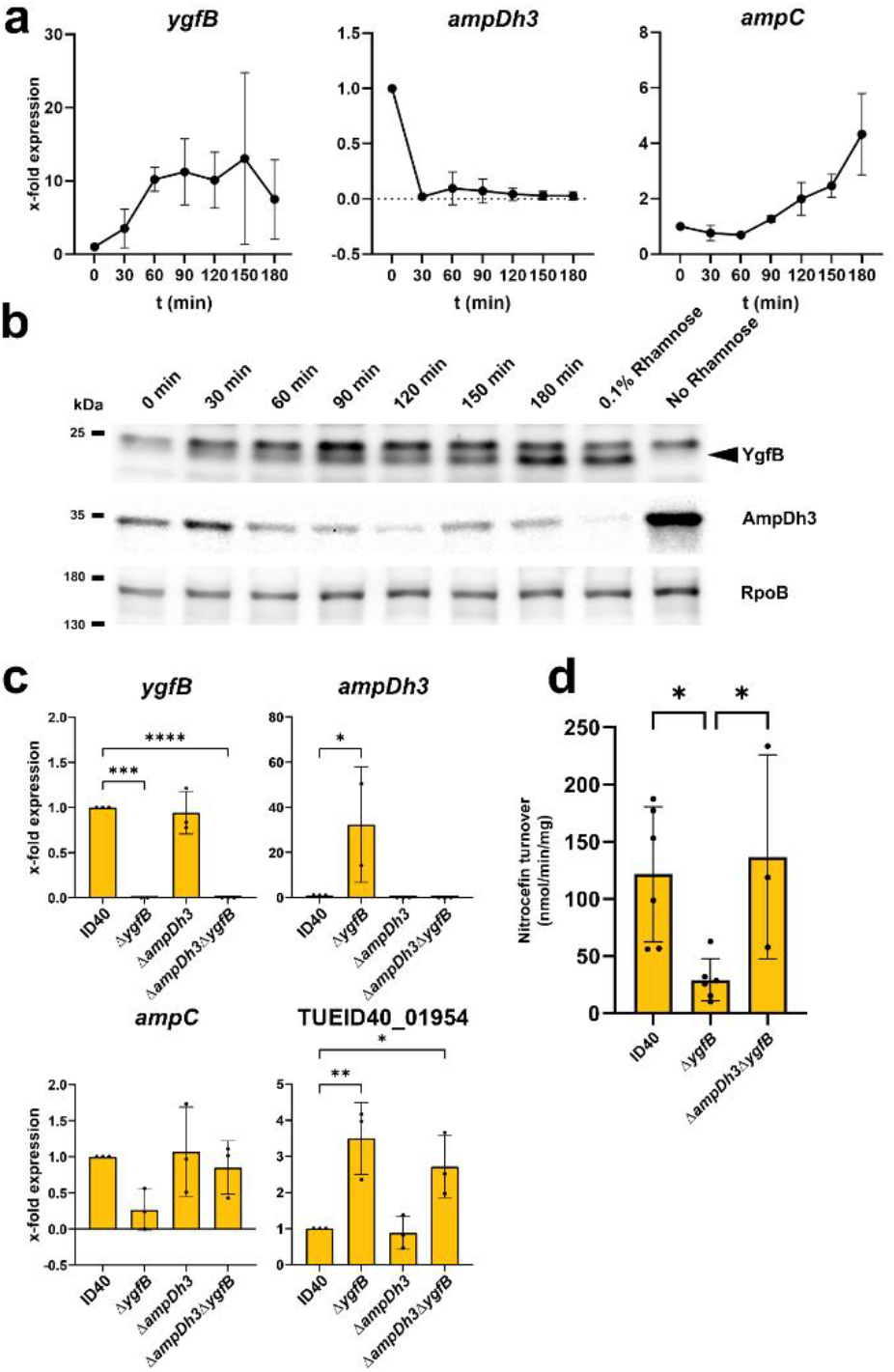
Relationship between YgfB, AmpDh3 and AmpC. (a) The expression of *ygfB* in ID40Δ*ygfB*::rha-*ygfB* was induced with 0.1% rhamnose at time point zero and RNA was isolated at various time points of growth in LB medium for further use in RT-qPCRs. Data depict the mean and SD of x-fold expression compared to ID40 0 min of *n*=2-3 independent experiments (b) The expression of *ygfB* in ID40Δ*ygfB*::rha-*ygfB*::*ampDh3*-HiBiT was induced with 0.1% rhamnose at time point zero and samples were taken from the growing culture in LB medium at the indicated time points. Then, whole cell lysates were prepared and used for SDS-PAGE and Western blots. The 0.1% condition depicts a strain grown under constant rhamnose supplementation. As primary antibodies, anti-YgfB or anti-RpoB antibodies and as secondary antibody anti-IgG-HRP antibodies were used and detection was done using ECL. For determination of AmpDh3, recombinant LgBiT was used. LgBiT binds to HiBiT resulting in a functional luciferase. The cleavage of the substrate furimazine leads to detectable chemiluminescence. Data are representative of three independent experiments. (c) *P. aeruginosa* strains as indicated were grown for 3 h in LB. RNA was isolated and used for RT-qPCR. Data depict mean and SD of x-fold mRNA expression compared to ID40 of *n*=2-3 independent experiments. (d) β-lactamase activity of indicated strains was determined by using a nitrocefin assay. Data depict mean and SD of *n*=3-6 independent experiments. Asterisks indicate significant differences compared to ID40 (*p<0.05, **p<0.01, ***p<0.001, ****p<0.0001; one-way ANOVA, Dunnett’s multiple comparisons comparing to ID40, (d) Šídák’s multiple comparisons)

### Deletion of YgfB leads to an AmpDh3-dependent reduction in AmpR-activating anhMurNAc-peptide levels

Next, we sought to understand the connection between AmpDh3 levels and *ampC* expression. According to the current literature, *ampC* expression is regulated by the transcriptional regulator AmpR in response to peptidoglycan turnover products and synthesis precursors^7^. AnhMurNAc-3P and anhMurNAc-5P are supposed to induce the activator function of AmpR, while UDP-MurNAc-5P induces the repressor function of AmpR in respect to AmpC β-lactamase expression^7, 8, 19^. From this we deduced that the deletion of *ygfB* might possibly indirectly change the composition of PG precursors, thereby affecting AmpC induction **(Fig. 3)**. HPLC-mass spectrometry analyses of cytosolic extracts revealed that *ygfB* deletion led to reduced levels of N-acetylglucosamine (GlcNAc)-anhMurNAc-3P as well as anhMurNAc-3P- and -5P, and increased levels of GlcNAc-anhMurNAc. No or only subtle changes in the levels of UDP-MurNAc-5P, UDP-MurNAc and anhMurNAc were found. Taken together, by inhibiting production of AmpDh3, YgfB reduces the degradation of anhMurNAc-peptides. Thereby, YgfB modulates the balance between anhMurNAc-peptides and UDP-MurNAc-5P, shifting it towards an anhMurNAc-peptide-response of AmpR; i.e., the upregulation of AmpC β-lactamase. From these data it can be concluded that YgfB-mediated repression of AmpDh3 production impacts the composition of PG precursors which in turn modulates *ampC* expression.

**Fig. 3:**
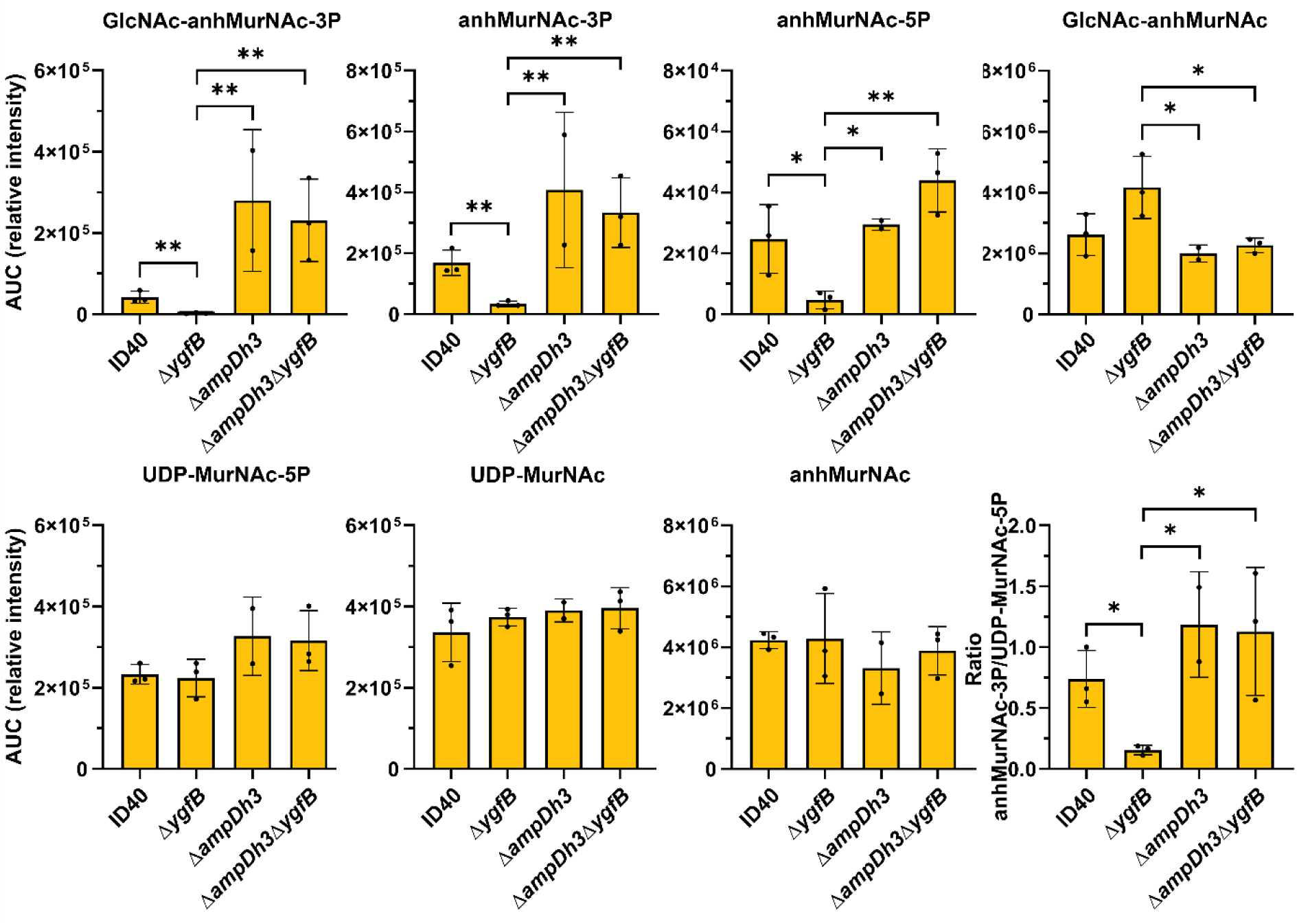
Impact of YgfB and AmpDh3 on the composition of peptidoglycan precursors. The indicated *P. aeruginosa* strains were grown for 6 h in LB medium, cytosolic extracts were generated and analyzed by HPLC-mass spectrometry. The graphs depict the mean and SD of the area under curve (AUC) of the peaks obtained for various catabolites of *n*=2-3 independent experiments. In addition, the ratio between anhMurNAc-3P and UDP-MurNAc-5P is shown. Asterisks indicate significant differences between *ΔygfB* and all other groups (*p<0.05, **p<0.01; Logarithmic values were shown to be normally distributed according to Wilks-test. Significance was then tested using Tukeys,,Honestly Significant Difference “-Test (*p<0.05, **p<0.01)

### AmpDh3 is located in the cytoplasm

Zhang et al. previously demonstrated that AmpDh3 and AmpDh2 preferentially cleave long-chain peptidoglycan fragments and have much lower activity toward anhydro-muramyl derivates^13^. This led to the assumption that both AmpDh2 and AmpDh3 have to be localized in the periplasm. We wanted to clarify where AmpDh3 is actually localized.

To assess the localization of AmpDh3, periplasmic proteins were separated from cytoplasmic proteins by spheroplasting and Western blot analysis was performed. While the periplasmic protein SurA was enriched in the periplasmic fraction, the cytoplasmic protein RpoB as well as AmpDh3 were found predominantly in the cytoplasmic fraction, indicating that AmpDh3 is mainly located in the cytoplasm **(Fig. S4).** Meanwhile, this was also confirmed by others^20^.

### YgfB represses AlpA-mediated AmpDh3 promoter activity

The transcriptome data suggested that YgfB suppresses *ampDh3* transcription. Previous studies described that both the *alpBCDE* cluster as well as the *ampDh3*-TUEID40_01954 (PA0808) operon are regulated by AlpA^16, 17^. AlpA seems to act as an antiterminator and to bind to the AlpA binding element (ABE) of both the *alpBCDE* genes, as well as the *ampDh3* promoter^16, 17^. Peña et al.^16^ defined a transcriptional start site (TSS) at position -413 bp upstream of the start of the coding sequence (CDS) of *ampDh3* and a minimal promoter region between positions -469 and -409 comprising an AlpA binding element **(Fig. 4a)**. In addition, they predicted two terminator regions starting at 26 bp downstream of the ABE and at bp -178 to -137 upstream of the *ampDh3* CDS. Among the two, the terminator region in close proximity to the ABE has only a minor impact on transcription^16^. In line with this, the terminator prediction tool ARNold^21^ suggests one terminator region between positions -178 to -137, corresponding to the second terminator region predicted by Peña et al.^16^.

**Fig. 4:**
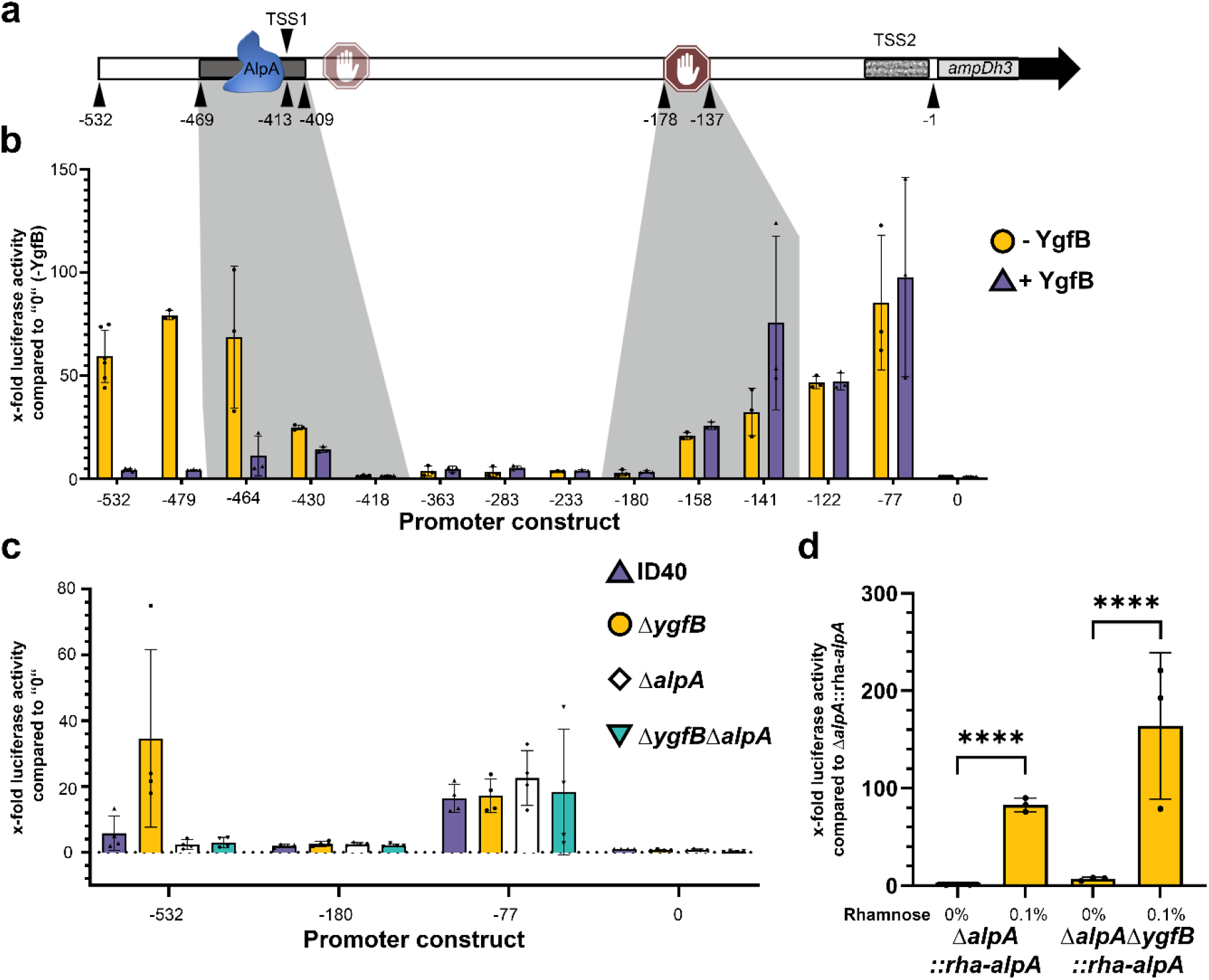
Analysis of the *ampDh3* promoter. (a) Schematic view of the putative *ampDh3* promoter. Numbers below depict the base pairs counted upstream of the coding sequence (CDS). The black bar in which blue AlpA is depicted marks the AlpA binding element (ABE) as defined by Peña et al^16^. Stop symbols mark terminator regions as predicted. (b and c) Luciferase assays were performed to measure transcriptional activity using *ampDh3* promoter fragments of various lengths fused to the coding sequence of NanoLuc. Data depict the mean and SD of the x-fold luciferase activity of 0-luc compared to all other fragments of *n*=3, *n*=6 for (b) and *n*=4 for (c) independent experiments. In (b), the strain ID40Δ*ygfB*::rha*-ygfB* without (yellow circle, depicted as -YgfB) or with 0.1% rhamnose (blue triangle, depicted as +YgfB) and in (c) the strains ID40, *ΔygfB, ΔalpA, ΔygfBΔalpA*, were used. The strains carry the indicated pBBR plasmids in which the expression of NanoLuc is under the control of fragments of different length of the *ampDh3* promoter. For instance, -532 is the abbreviation for a pBBR plasmid containing an *ampDh3* promoter fragment comprising the region -532 to -1 bp upstream of the CDS of NanoLuc. In (d) the strains Δ*alpA*::rha-*alpA* and *ΔygfB ΔalpA::rha-alpA* carrying pBBR-532-luc were used as indicated. Asterisks in (d) indicate significant differences (*p<0.05, **p<0.01, ***p<0.001, ****p<0.0001; one-way ANOVA, Šídák’s multiple comparisons)

To assess the impact of YgfB on the activity of the *ampDh3* promoter, various reporter constructs were generated. This was done by fusing *ampDh3-*promoter fragments of various lengths (starting between bp -532 to bp -77 and ending at bp -1) to a reporter gene encoding the luciferase NanoLuc, using the vector pBBR1. To assess background activity, NanoLuc was cloned into pBBR1 without any promoter (designated in **Fig. 4b** as 0). These fragments were introduced into the conditional *ygfB* mutant ID40Δ*ygfB*::rha-*ygfB*. Subsequently, luciferase assays were performed both in the presence and in the absence of rhamnose to determine the transcriptional activity of the *ampDh3* promoter in a YgfB-dependent manner **(Fig. 4b).**

Interestingly, a promoter fragment comprising bp -77 to bp -1 was sufficient to observe YgfB-independent promoter activity. A promoter fragment comprising bp -180 to bp -1 abrogated nearly all promoter activity, which is in line with the findings of Peña et al.^16^, and the existence of a terminator region. The shortest promoter fragment showing YgfB-dependent transcriptional regulation of *ampDh3* was the fragment between positions -464 and -1 comprising almost the entire suggested ABE. These data indicate that transcription can be initiated either at position -413 upstream of the *ampDh3* CDS (here referred as transcription start site 1, TSS1) or upstream of position -77 (transcription start site 2, TSS2). However, transactivation at TSS2 is not regulated by YgfB. YgfB-dependent inhibition of *ampDh3* transactivation seems to occur in the same region in which AlpA binds.

Next, the luciferase reporter plasmids 0, -77, -180 and -532 were introduced into the strains ID40, ID40Δ*ygfB*, ID40Δ*alpA* and ID40Δ*alpA*Δ*ygfB*, as well as 0 and -532 into ID40Δ*alpA*::rha-*alpA* and ID40Δ*ygfB*Δ*alpA*::rha-*alpA* and again the luciferase activity was measured as a proxy for the promoter activity **(Fig. 4c and Fig. 4d)**. We found that the activity of the longest promoter fragment (−532) increased upon deletion of *ygfB*. Additional deletion of *alpA* abrogated promoter activity demonstrating that AlpA is the essential factor to trigger *ampDh3* transactivation, while YgfB counteracts AlpA-induced promoter activity (**Fig. 4c**). Complementation experiments (**Fig. 4d**) show that the induction of AlpA with rhamnose results in high *ampDh3* promoter activity. The promoter fragment -180, including the terminator region, revealed no promoter activity in any of the strains (**Fig. 4c**). However, the promoter fragment -77 showed promoter activity independent of the presence of either AlpA or YgfB (**Fig. 4c**). Hence, this confirms our results obtained with the conditional *ygfB* mutant ID40Δ*ygfB*::rha-*ygfB* and suggests an interplay between AlpA and YgfB.

### A functional PBP4 is required for basal expression of AmpDh3

Previous studies using the strain PAO1 revealed that the deletion or mutation of *dacB*, encoding the D-Ala-D-Ala-carboxypeptidase PBP4, leads to an overproduction of AmpC and results in hyperresistance^5^. Of note, the MDR *P. aeruginosa* strain ID40 carries a mutation that inactivates the *dacB* gene^10, 11^. Furthermore, it was suggested that the activity of the two-component system CreBC is triggered by *dacB* inactivation in response to β-lactams and affects antibiotic resistance^5^. Thus, we asked whether *dacB* inactivation and the subsequent activation of the two-component system CreBC may also influence the expression of *ampDh3* and therefore affect β-lactam resistance. For this purpose, we first investigated whether *dacB, creBC* or *ampR* inactivation contributes to the repression of *ampDh3* transactivation. To achieve this, an *ampDh3*-532-luc reporter construct was introduced into various mutants. Neither the deletion of *ampR* nor the deletion of *creBC* had an impact on the *ampDh3* promoter activity **(Fig. 5a).** However, complementation of ID40 (naturally equipped with a non-functional *dacB)* with a functional *dacB* derived from PA14 led to a two-fold increase in *ampDh3* promoter activity after induction with rhamnose **(Fig. 5b)**. This finding indicates that PBP4 inactivation suppresses the transactivation of the *ampDh3* promoter to some extent. The impact of a non-functional *dacB* gene on *ampDh3* expression levels, however, is low in comparison to the impact of *ygfB* deletion. This suggests that the YgfB-dependent repression of *ampDh3* is largely independent from the effects of a loss of PBP4 function.

**Fig. 5:**
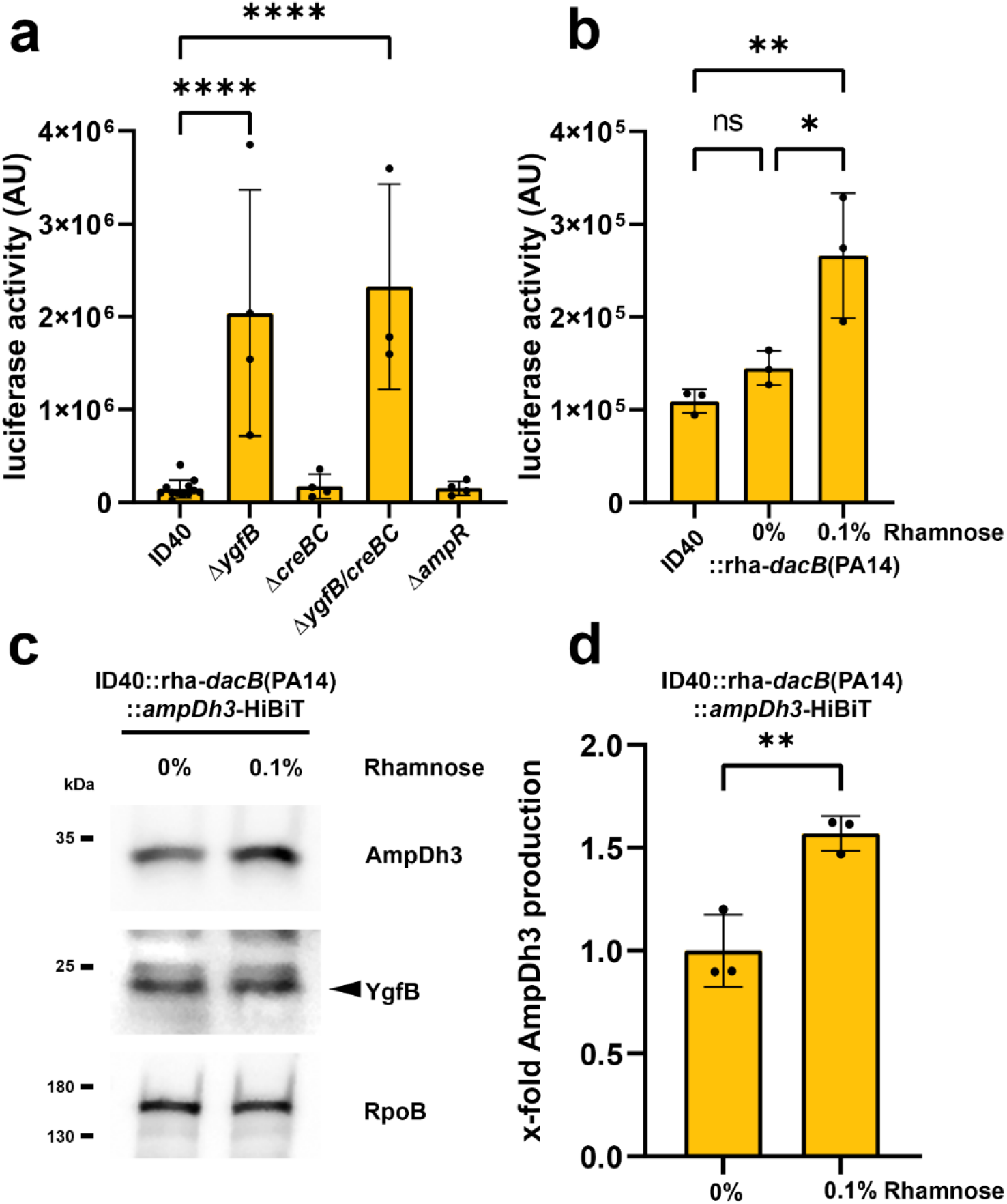
Interrelationship between *creBC, dacB* and *ampDh3*. (a and b) Luciferase reporter assays using the indicated strains harboring the reporter construct *ampDh3*-532-luc comprising the *ampDh3* promoter fragment between position -532 and -1 upstream of the CDS of Nanoluc. Data depict the mean and SD of luciferase activity of *n*=12, 4, 4, 3, 4 for (a) and for (b) *n*=3 independent experiments. Asterisks in (a) indicate significant differences compared to ID40 (****p<0.0001; one-way ANOVA, Dunnett’s multiple comparisons comparing to ID40) and in (b) compared between all conditions (ns p>0.05, *p<0.05, **p<0.01; one-way ANOVA, Tukey’s multiple comparisons). (c) ID40::rha-*dacB*(PA14)::*ampDh3*-HiBiT was grown for 3 h without or with 0.1% rhamnose to induce the expression of *dacB* of PA14 in LB medium at 37°C. Whole cell lysates were used to perform SDS-PAGE and Western blot transfer. As primary antibodies anti-YgfB or anti-RpoB antibodies and as secondary antibodies anti-IgG-HRP antibodies were used and detection was done using ECL. AmpDh3-HiBiT was detected by incubating the membrane with LgBiT and furimazine. Data are representative for three independent experiments. (d) Bacteria were harvested, lysed and subsequently luciferase activity of AmpDh3-HiBiT was measured as described in material and methods. Data depict the mean and SD of the luciferase activity of *n*=3 independent experiments. Asterisks indicate significant differences (p**<0.01; unpaired, two-tailed t-test).

Nevertheless, a functional PBP4 shows a higher basal transcriptional activity of the *ampDh3* promoter. To further check this, AmpDh3 production was investigated in the strain ID40::rha-*dacB*(PA14)::*ampDh3*-HiBiT. Our results show that upon induction of *dacB* expression, the AmpDh3 production was indeed ~1.5 times higher compared to uninduced cells **(Fig. 5c and d)**.

### Repression of AlpA-mediated AmpDh3 production by YgfB prevents higher levels of susceptibility to β-lactam antibiotics as well as to the combination of β-lactam antibiotics and ciprofloxacin

As previously shown, *ygfB* deletion increases the susceptibility of ID40 to β-lactam antibiotics. To investigate the relationship between YgfB and AlpA/AmpDh3, the minimum inhibitory concentration (MIC) of several β-lactam antibiotics was measured in various mutants. As indicated in **Fig. 6**, deletion of *ygfB* led to increased susceptibility to all tested antibiotics. Additional deletion of either *alpA* or *ampDh3* restored most of the MIC values to those of ID40 WT. These data indicate that in ID40, YgfB contributes to resistance to β-lactam antibiotics by repressing AlpA-mediated AmpDh3 production.

**Fig. 6:**
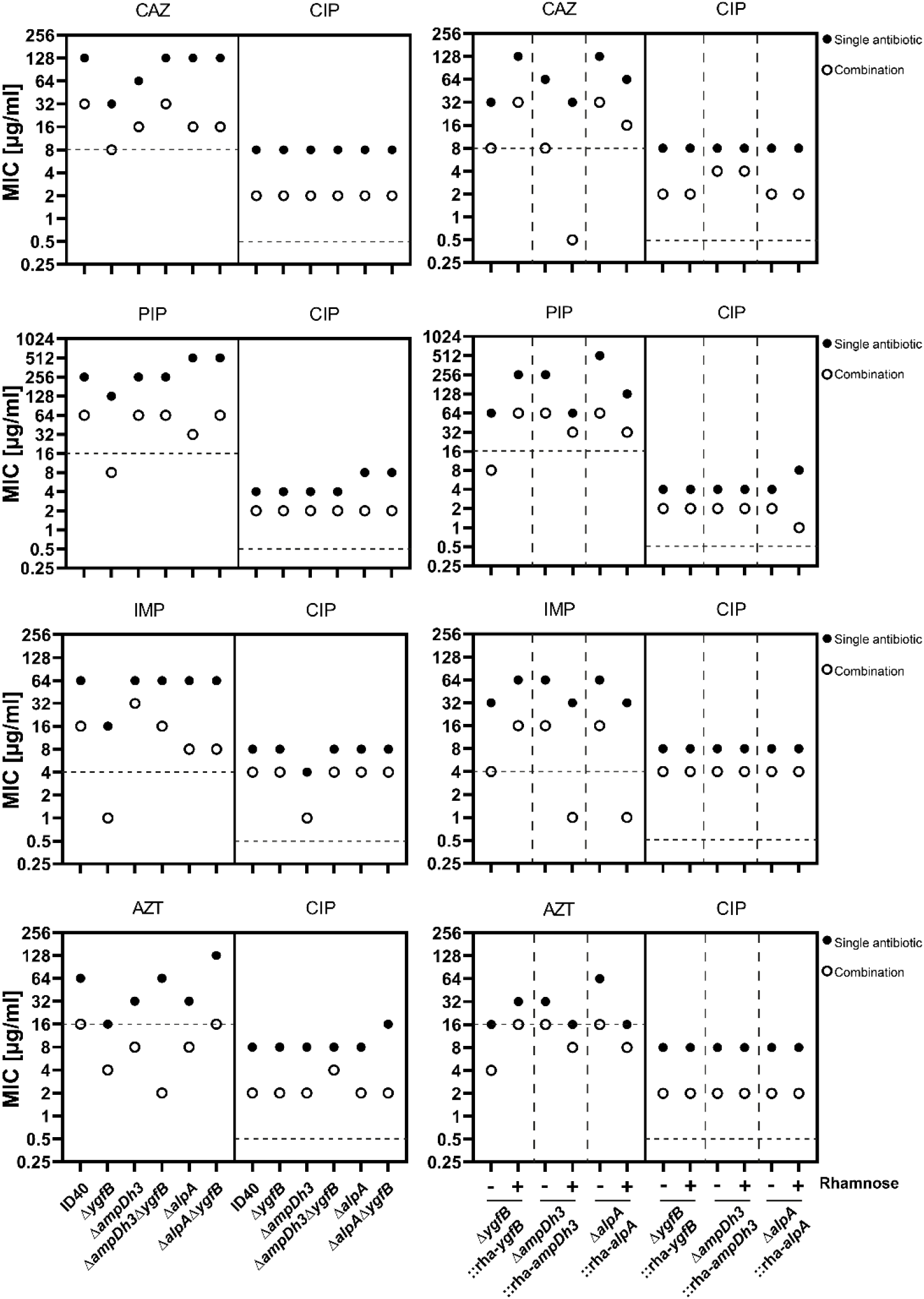
Checkerboard assays to investigate combinatory effects of ciprofloxacin and β-lactams. The indicated antibiotics were combined in log2-fold dilutions and minimum inhibitory concentrations for antibiotics alone or in combination were determined. Plotted are the MICs for single antibiotics (filled circle) and combination of antibiotics (open circle). The left panel shows single deletion mutants and the right panel shows conditional deletion mutants grown in the absence or presence of 0.1% rhamnose. Dotted horizontal lines show break points for each antibiotic according to EUCAST. CAZ ceftazidime, PIP piperacillin, AZT aztreonam, IMP imipenem, CIP ciprofloxacin. Data are derived from two independent experiments for each combination.

DNA damage by ciprofloxacin was shown to lead to autocleavage of the repressor AlpR, which controls *alpA* expression. In turn, this results in increased AlpA-mediated transactivation of the *ampDh3* promoter^15, 16^. We hypothesized that a ciprofloxacin-mediated increase in AmpDh3 production might increase the susceptibility of ID40 or ID40Δ*ygfB* toward β-lactam antibiotics. To investigate this, checkerboard analyses were performed by combining various β-lactam antibiotics with ciprofloxacin to determine MIC values (**Fig. 6**) as well as fractional inhibitory concentration indices (FIC-I) (**Table S1**). FIC-I is a measurement of the combined effect of an antibiotic combination. For experimental details and calculation of FIC-I please refer to the material and methods section under checkerboard assays. In all tested strains MIC values for ciprofloxacin varied between 4-8 μg/ml and in combination with the tested β-lactam antibiotics the MIC value ranged mostly between 2-4 μg/ml.

Measurement of MIC values for ID40, *ΔampDh3* or *ΔalpA* showed that the combination of CIP and β-lactam antibiotics reduced MIC values for the tested antibiotics in all three strains compared to single treatment in a similar manner. Deletion of *ygfB* further decreased the MIC values by two to three log2 steps as compared to ID40 depending on the β-lactam antibiotic used. This can be explained by the AlpA-induced AmpDh3 production, as additional deletion of either *alpA* or *ampDh3* restored the resistance to β-lactam antibiotics to the levels of ID40 wildtype. Additionally, experiments were performed using conditional mutants (Δ*ygfB*::rha-*ygfB*, Δ*alpA*::rha-*alpA*, *ΔampDh3::rha-ampDh3)* in which the gene of interest can be induced with rhamnose **(Fig. 6 right column)**. In the absence of rhamnose, the conditional deletion mutants responded to treatment with various β-lactams, alone or in combination with ciprofloxacin, like the corresponding single deletion mutants as confirmed by comparison of the MIC values.

Upon rhamnose supplementation the MIC values of antibiotics measured for the conditional *ygfB* deletion mutant were similar to the MIC values for the ID40 wildtype. In contrast, the conditional *ampDh3* and *alpA* deletion mutants showed lower β-lactam resistance upon rhamnose treatment. This indicates, that under these conditions, the effect of YgfB can be overridden.

The fractional inhibitory concentration index (FIC-I) was calculated for the checkerboard analyses to distinguish between synergistic and additive effects of the antibiotic combinations **(Table S1)**. Regarding all tested antibiotics and strains, the FIC-I ranged between 0.25 to 0.75 (with one exception: FIC-I 1 for the conditional *ampDh3* deletion mutant treated with rhamnose using the PIP/CIP combination). Following the definition that FIC-I values ≤0.5 indicate synergism, while values between >0.5 and 1 indicate additive effects, both synergistic and additive effects can be observed (see also discussion).

To monitor AmpDh3 production 18 h after treatment with or without 2.5 μg/ml ciprofloxacin, AmpDh3-HiBIT production in the strains ID40::*ampDh3-*HiBiT and *ΔygfB::ampDh3-HiBiT* was investigated. In ID40::*ampDh3-*HiBiT, only very low levels of AmpDh3 were detected, but they were increased by the deletion of *ygfB* **(Fig. S5)**. Ciprofloxacin treatment and even more so, *ygfB* deletion strongly increased the AmpDh3 production.

Taken together, ciprofloxacin and β-lactam antibiotics have a rather additive than synergistic inhibitory effect on the investigated ID40 strain. The repressive effect of YgfB on AlpA-mediated AmpDh3 production prevents that in combination with ciprofloxacin, resistance of ID40 to β-lactam antibiotics can be broken.

### YgfB interacts with AlpA and interferes with AlpA-DNA binding

The repressive action of YgfB on the *ampDh3* cluster and to some extent the *alpBCDE* cluster led to the idea that YgfB may directly act on either AlpR or AlpA. To address this, the strains ID40::HA-*alpR*::*alpA*-HiBiT and *ID40ΔygfB::HA-alpR::alpA-HiBiT* were grown for two hours without or with 32 μg/ml ciprofloxacin and Western blot analyses were performed **(Fig. 7).**

**Fig. 7:**
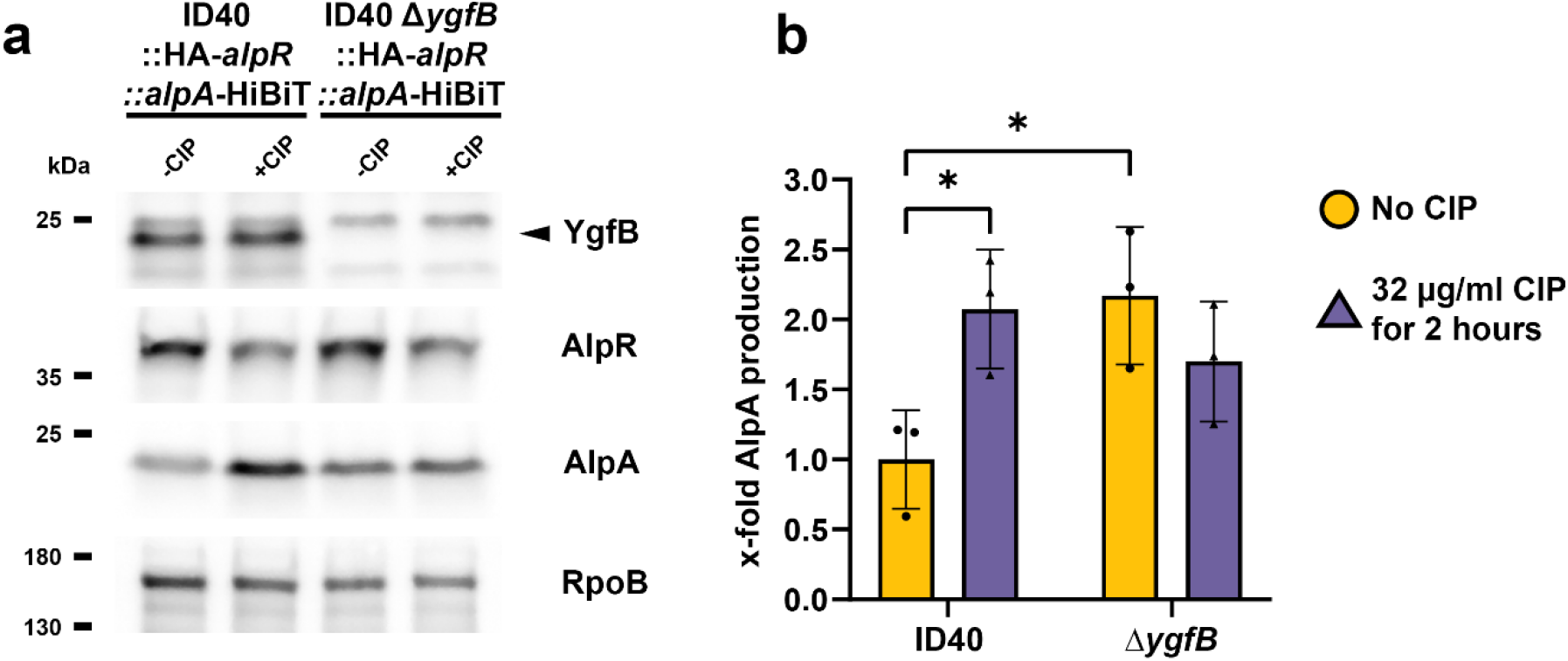
Modulation of AlpR and AlpA production by YgfB and ciprofloxacin (CIP). (a) Whole cell lysates of the indicated strains were used for Western blot analyses. +CIP conditions were treated with 32 μg/ml for 2 hours. As primary antibodies, anti-HA, anti-YgfB or anti-RpoB antibodies and as secondary antibody anti-IgG-HRP antibodies were used and detection was done using ECL. For detection of AlpA-HiBiT or AmpDh3-HiBiT, recombinant LgBiT was used. LgBiT binds to HiBiT resulting in a functional luciferase. The cleavage of the substrate furimazine leads to detectable chemiluminescence. Data are representative of three independent experiments. (b) Quantification of AlpA by measuring luciferase activity of AlpA-HiBiT in lysed cell extracts. Data depict mean and SD of x-fold luciferase activity compared to ID40 WT for *n*=3 independent experiments. Asterisks indicate significant differences compared to ID40 -CIP (*p<0.05; two-way ANOVA, Šídák’s multiple comparisons comparing to ID40 -CIP).

We observed that only ciprofloxacin but not the deletion of *ygfB* reduced the levels of AlpR. Following ciprofloxacin treatment, the levels of AlpA were upregulated in ID40. However, while deletion of *ygfB* increased AlpA levels when compared to ID40, combining the deletion and ciprofloxacin treatment had no further effect on the protein levels (**Fig. 7a**).

To confirm the Western blot results, we additionally analyzed the levels of AlpA-HiBiT using the Nano-Glo HiBiT Lytic Detection System (Promega). We observed increased AlpA-HiBiT protein levels (1.6 to 2-fold) in the *ygfB* deletion mutant as assessed by measuring the luciferase activity of the fusion protein (**Fig. 7b**). Ciprofloxacin increased the levels in a similar fashion, but once again we did not observe a synergistic effect on AlpA levels upon concurrent ciprofloxacin treatment and *ygfB* deletion.

Since YgfB seemed to influence AlpA but not AlpR, we investigated whether AlpA interacts with YgfB (**Fig. 8**). In a first attempt, recombinant GST-YgfB or GST was mixed with cell lysates of the ID40Δ*ygfB*::HA-*alpR*::*alpA*-HiBiT strain and a GST pull down assay was performed **(Fig. 8a)**. In the subsequent Western blot analysis, we could detect a band for AlpA-HiBiT when the lysates were mixed with GST-YgfB but not when mixed with GST, indicating that YgfB does interact with AlpA. To clarify whether the interaction between YgfB and AlpA is direct or indirect, we performed pulldown assays with recombinant His-MBP-AlpA or His-MBP as a bait and recombinant YgfB as a prey **(Fig. 8b)**. We observed an enrichment of YgfB when incubated with His-MBP-AlpA. This pointed to a direct interaction between AlpA and YgfB and led us to the hypothesis that the interaction of YgfB with AlpA might abrogate the ability of AlpA to bind to the *ampDh3* promoter. To test the binding of AlpA to its binding site in the *ampDh3* promoter region, electrophoretic mobility shift assays (EMSAs) were performed using a probe comprising the ABE element. Addition of His-MBP-AlpA, but not the addition of His-MBP resulted in a weak mobility shift of the fluorophore-labeled DNA probe, with the band appearing smeary **(Fig S6)**. Numerous attempts to optimize the procedure failed, but this might be attributed simply to the nature of this interaction which is presumably rather weak. While the addition of bovine serum albumin to His-MBP-AlpA did not affect the formation of a complex between AlpA and the ABE, the addition of recombinant YgfB led to a decrease in the formation of the AlpA-ABE complex to a level resembling the amount of His-MBP-AlpA bound non-specifically to a scrambled DNA probe. **(Fig. S6)**.

**Fig. 8:**
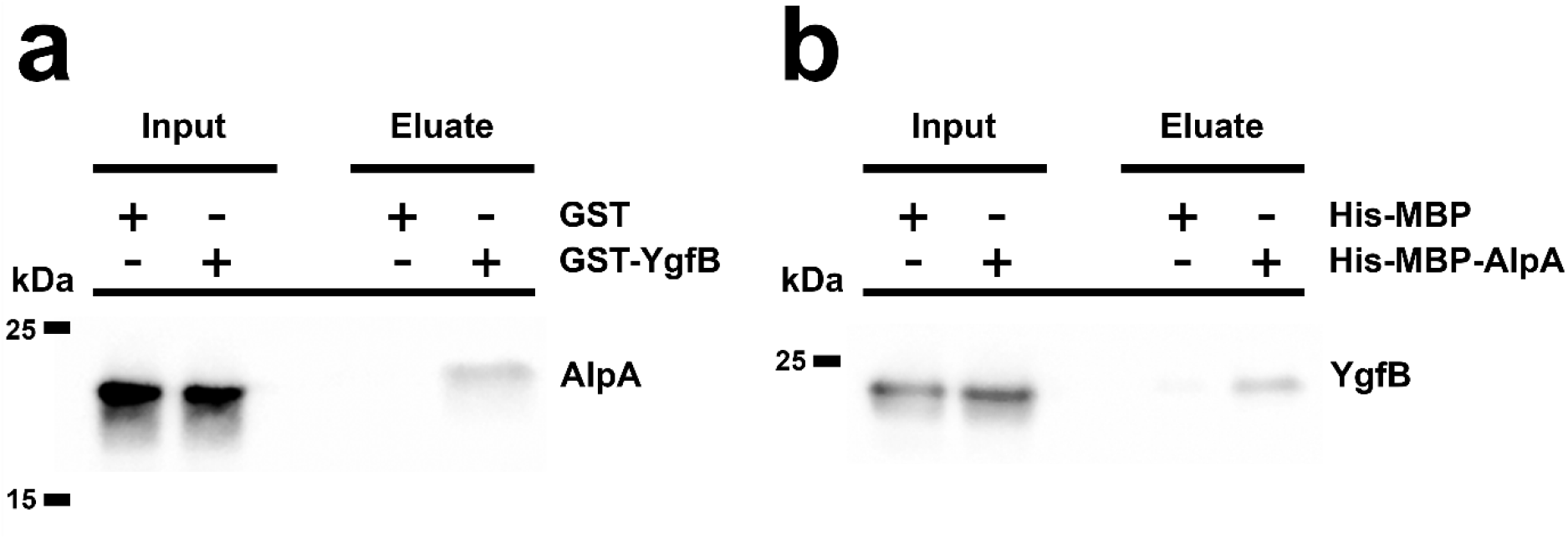
Interaction of AlpA with YgfB. Pulldowns were performed using (a) recombinant GST-YgfB or GST as a bait and cell lysates of ID40Δ*ygfB*::*alpA*-HiBiT::HA-*alpR* as the prey and (b) recombinant His-MBP or His-MBP-AlpA as a bait and recombinant YgfB as the prey. Shown are Western blots using either LgBiT or anti-YgfB antibodies for detection. Of note: the slight difference in migration of AlpA/YgfB in input and eluate is associated with the different composition of input and elution buffer (a) is representative for five experiments, (b) for two experiments.

We are aware that the low quality of the EMSA limits its informative value to some extent. However, we think that these results, especially if considered in conjunction with the results presented before, strongly suggest that the interaction of YgfB with AlpA most likely abrogates the binding of AlpA to the ABE. This finding is consistent with the inhibitory effect of YgfB on AlpA-mediated transactivation.

### The role of YgfB in β-lactam resistance is conserved in other *P. aeruginosa* strains

To investigate whether the prominent role of YgfB in β-lactam resistance holds true also for other MDR *P. aeruginosa* strains, *ygfB* was deleted in the clinical blood stream infection isolates ID143 and ID72 as well as in the more sensitive strains PAO1 and PA14. As depicted in **Table S2**, β-lactam resistance was also decreased in the investigated *P. aeruginosa* strains, indicating that the presence of YgfB seems to be of general importance to achieve higher resistance of *P. aeruginosa* to β-lactam antibiotics. In addition, in all tested strains, the deletion of *ygfB* led to higher *ampDh3* promoter activity **(Fig. S7a)**. However, regarding the tested strains so far it cannot be stated that lower levels of basal *ampDh3* promoter activity are a specific characteristic of more resistant strains, since the MDR strain ID40 and the sensitive strain PAO1 show similar basal *ampDh3* promoter activity **(Fig. S7b)**.

## Discussion

We have previously demonstrated that the deletion of *ygfB* reduces resistance against β-lactam antibiotics in an MDR *Pseudomonas aeruginosa* strain^10^. In the present study the function of YgfB in relation to β-lactam resistance was addressed. Our findings revealed that (i) YgfB-mediated suppression of the AlpA-induced production of AmpDh3 contributes to β-lactam resistance. Lack of YgfB would otherwise lead to higher AmpDh3 levels, which change the composition of peptidoglycan turnover products resulting in the AmpR-mediated reduction of *ampC* expression. (ii) YgfB interferes with the transactivating function of AlpA by directly interacting with AlpA and likely preventing it from binding to the AlpA binding element. (iii) Ciprofloxacin-induced DNA damage increases the susceptibility of *P. aeruginosa* to β-lactam antibiotics, an effect that is antagonized by the YgfB-mediated negative regulation of AmpDh3. (iv) Inactivation of *dacB* dampens the transcriptional activity of the *ampDh3* promoter to some extent. This may be an additional aspect of how *dacB* inactivation leads to higher resistance to β-lactam antibiotics.

Comparative transcriptome analyses of ID40 and ID40Δ*ygfB* revealed very few transcriptional changes, including the downregulation of *ampC* and the upregulation of the *ampDh3-TUEID40_01954* operon and the *alpBCDE* cluster. The *alpBCDE* cluster was recently described to be a potential self-lysis mechanism important for successful lung infection by *P. aeruginosa*^15^. The effect of YgfB on cell death induction is so far unclear and requires follow-up studies.

*P. aeruginosa* possesses three paralogous amidases called AmpD, AmpDh2 and AmpDh3^13, 22^. It was already shown that loss of function mutations in *ampD* are a frequent source of MDR^4, 7^. In addition, a deletion of only *ampD* and, to a lesser extent, of either *ampDh2* or *ampDh3* was shown to increase the resistance of PAO1 to β-lactam antibiotics^23^. However, deleting both *ampD* and *ampDh3* leads to a much more pronounced resistance which can hardly be increased by additional deletion of *ampDh2*^23^. The increasing resistance associated with one or several of these amidases is also correlated with increasing levels of the cephalosporinase AmpC^23^.

As demonstrated in the presented study AmDh3 seems to act redundantly with AmpD and localizes to the cytoplasm as also shown by Colautti et al^20^.

Our data clearly indicate that YgfB increases *ampC* expression in an AmpDh3-dependent manner by decreasing the levels of the AmpR-activating anhMurNAc-peptides^7, 8, 19^. The balance between the AmpR-inactivating UDP-MurNAc-peptides and the AmpR-activating anhMurNAc-peptides can be shifted to promote the activator function of AmpR by additional deletion of *ampDh3*. Interestingly, in the single *ygfB* deletion mutant the reduced levels of anhMurNAc-3P and -5P do not result in higher levels of anhMurNAc. However, in the *ygfB* deletion mutant the levels of N-Acetylglucosamine (GlcNAc)-anhMurNAc-3P were decreased and the GlcNAc-anhMurNAc levels were increased. This suggests that the main target of AmpDh3 might be primarily the GlcNAc-anhMurNAc-peptides.

Thus, all these data, including the determination of the MIC values clearly demonstrate that YgfB contributes to β-lactam resistance by repressing *ampDh3* expression and thereby greatly altering the anhMurNAc-peptide/UDP-MurNAc-peptide balance and the level of *ampC* expression. Changes in *ampDh3* promoter activity and antibiotic resistance due to the deletion of *ygfB* seem not to be limited to the ID40 strain, but could also be demonstrated for other MDR *P. aeruginosa* strains. However, the basal activity of AmpDh3 production does not seem to be directly linked to antibiotic resistance, meaning that the basal transcriptional activity of the *ampDh3* promoter cannot be used as a marker for sensitive or resistant strains.

Investigation of the role of *dacB* in YgfB-mediated suppression of *ampDh3* expression revealed that *dacB* inactivation slightly suppresses AmpDh3 production in a so far unknown and YgfB-independent manner. It can be concluded that the combined impact of YgfB and *dacB* inactivation on AmpDh3 suppression seems to be an important determinant of MDR in ID40.

A recent report showed that AlpA not only transactivates the *alpBCDE* cluster but also the *ampDh3* operon^16^. The authors proposed that AlpA binds to the ABE on the *ampDh3* promoter and then to the RNA polymerase. This binding allows the polymerase to bypass downstream terminators, resulting in *ampDh3* transcription^16^. As demonstrated in this study, deletion of *alpA* abrogates *ampDh3* transactivation of a complete *ampDh3* promoter. Our investigations confirmed that for AlpA-positive as well as YgfB-negative regulation of the *ampDh3* promoter the same upstream stretch including the ABE is very likely required. In addition, the importance of the second terminator region could be confirmed by showing that a promoter fragment comprising bp -180 to -1 upstream of the CDS does not show transcriptional activity. However, a fragment comprising bp -77 to -1 upstream of the CDS is sufficient for high transactivation in an AlpA/YgfB-independent manner. *In silico* analysis did not reveal a putative promoter element and at this point the second transcription initiation site is hypothetical and has to be defined in more detail.

As AlpA and YgfB only impacted the *alpBCDE* and the *ampDh3* operon we hypothesized that YgfB somehow interferes directly with the AlpR-AlpA axis. The model proposed suggests that AlpA first binds to the promoter at the putative ABE, and then to the RNA polymerase (RNAP), allowing RNAP to bypass the intrinsic terminator positioned downstream^16, 17^. Recently, Wen et al. confirmed these data by solving the AlpA-loading complex consisting of a nucleic acid scaffold corresponding to the positions -31 to 31 of the P_*alpB*_ promoter together with RNAP, σ^70^ and AlpA by cryo-EM^17^. These data might suggest, that for a robust binding of AlpA to the ABE, stabilization by RNAP and σ^70^ seems to be required. In contrast to Wen et al., we did not succeed to obtain native AlpA and therefore used a His-MBP tag to solubilize AlpA. We speculate that the addition of the His-MBP tag to the AlpA in combination with the lack of the other components of the AlpA-loading complex such as RNAP and σ^70^ is very likely the reason why we ended up with a highly reproducible but only weak binding of AlpA to the ABE. Nevertheless, this weak binding of AlpA was reliably abrogated by the addition of YgfB but not BSA that was used as negative control.

Taking also into account that YgfB directly interacts with AlpA and interferes with *ampDh3* transactivation in the region where the ABE is located, we think it is reasonable with all given caution to draw the conclusion that the direct interaction of YgfB with AlpA interferes with the binding of AlpA to the ABE and consequently with AlpA-mediated transactivation.

Several studies have described a synergism between ciprofloxacin and β-lactam antibiotics such as ceftazidime or aztreonam. For instance, Bosso et al. observed that in a total of 96 investigated *P. aeruginosa* strains, approximately 30% showed a synergistic inhibitory effect of ciprofloxacin and aztreonam or ceftazidime^24^. For *P. aeruginosa* strains which were both aztreonam and ciprofloxacin resistant, synergy was observed in 41% and resistance was broken for both antibiotics in 29% of such strains. When combining ciprofloxacin and ceftazidime, synergy was observed for 67% of the CIP/CAZ resistant strains, however, for none of the *P. aeruginosa* strains resistance to both antibiotics could be broken. Various other studies reported similar results, namely, that synergy between β-lactam antibiotics and ciprofloxacin can be observed for some but not all the strains of *P. aeruginosa*^25, 26, 27^. ID40 is a strain resistant to CIP and most β-lactam antibiotics. The definition of synergism by interpretation of fractional inhibitory concentration index (FIC-I) has been the subject of debate. While some publications interpret an FIC-I of <1 as synergistic^28, 29^ we have defined the interpretation as follows in accordance with several other sources^30, 31, 32^. A FIC-I of ≤0.5 indicates synergistic effects, an index between >0.5 and 1 additive effects. Indifferent effects are observed for values between >1 and 4 and values >4 indicate antagonistic effects.

The MIC values for ciprofloxacin varied randomly between 4 and 8 μg/ml in the performed checkerboards. This variation might influence the outcome of the presented FIC values and their interpretation. In the checkerboard assays performed for ceftazidime and aztreonam, the single MIC values for ciprofloxacin were 8 μg/ml with FIC-I values for ID40 and most of the deletion mutants smaller or equal to 0.5, indicating synergism. In contrast, in the checkerboard assays performed for piperacillin and imipenem, the MIC-values for ciprofloxacin had more variation, fluctuating between 4 and 8 μg/ml. The FIC-I was sometimes lower but mostly higher than 0.5 for the tested strains, indicating rather additive effects. Thus, a clear statement whether the combination of ciprofloxacin and β-lactam antibiotics is synergistic or additive in ID40 cannot be made. However, for the interpretation of the data on the effect of YgfB, the impact of such a distinction seems to be negligible.

The decisive point is that the repressive effect of YgfB on AlpA-mediated AmpDh3 production prevents that resistance can be broken for all tested β-lactam antibiotics upon additional treatment with CIP. Moreover, the action of YgfB might explain why CIP/β-lactam combinations are insufficient to kill many *P. aeruginosa* strains and highlights YgfB as an important contributor to β-lactam resistance as summarized in **Fig. 9.**

**Fig. 9:**
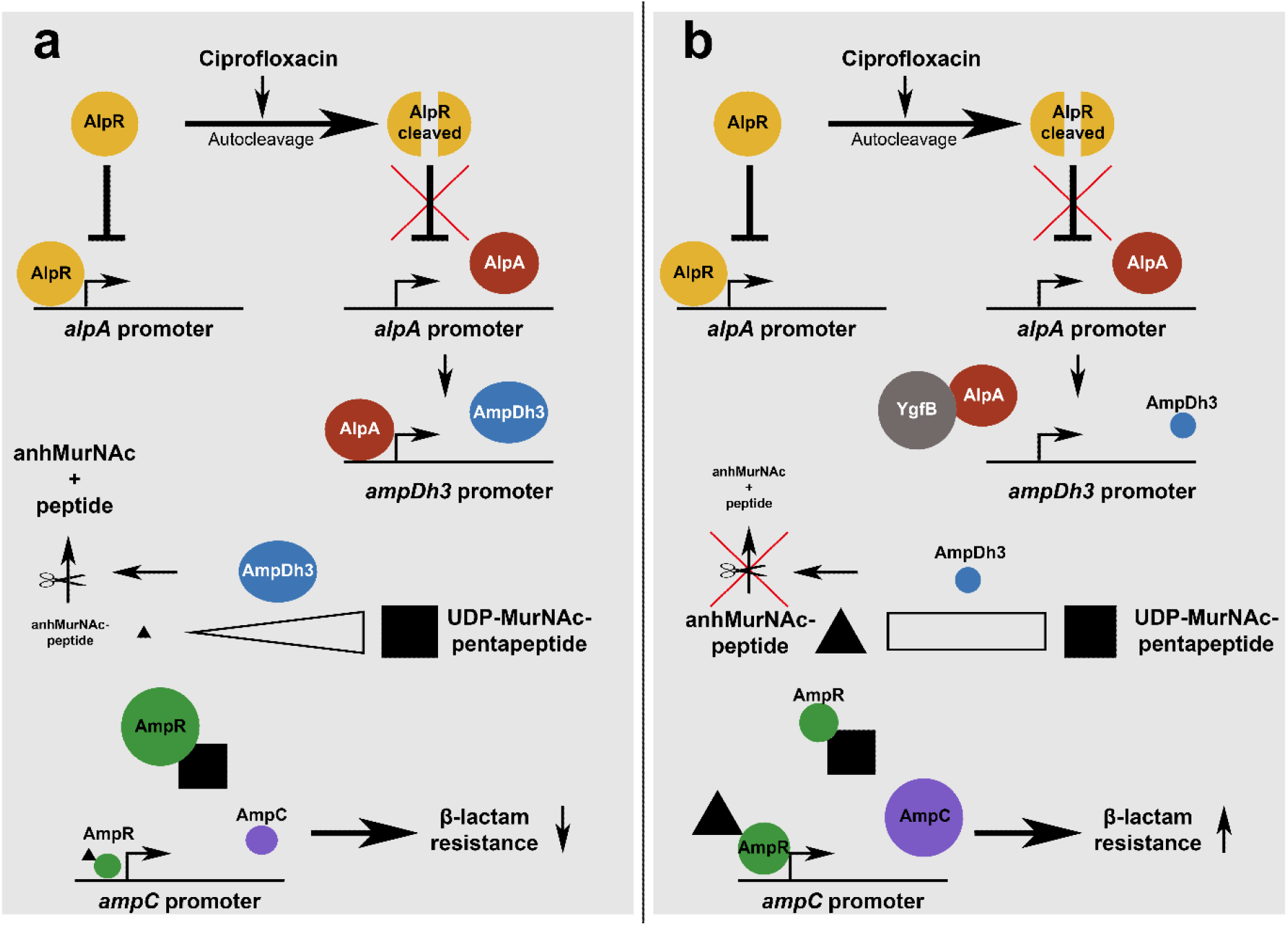
Ciprofloxacin-induced signaling pathway leading to modulation of *ampC* expression. (a) Ciprofloxacin triggers DNA damage which promotes autocleavage of AlpR and as a consequence, the derepression of the *alpA* promoter. AlpA acts as an antiterminator of *ampDh3* expression increasing AmpDh3 production. AmpDh3 cleaves anhMurNAc peptides which changes the balance of the AmpR-activator upon binding anhMurNAc-peptides and the AmpR-repressor upon binding UDP-MurNAc-5P in favor of AmpR-mediated repression of its target promoters. This leads to reduced *ampC* expression and decreased β-lactam resistance. (b) YgfB directly interacts with AlpA and blocks binding of AlpA to the *ampDh3* promoter. Thereby, AlpA’s antiterminator function is inhibited and *ampDh3* expression dampened. Reduced levels of AmpDh3 in the cytosol change the balance of AmpR-activating anhMurNAc-peptides and AmpR-repressing UDP-MurNAc-5P in favor of AmpR-mediated activation of its target promoters. This enhances AmpC production and leads to increased β-lactam resistance.

## Material and Methods

### Bacterial strains and culture conditions

Bacterial strains and plasmids used in this study are listed in **Supporting Table S3.** Bacteria were cultivated overnight at 37°C with shaking at 200 rpm in lysogeny broth (LB) containing suitable antibiotics if necessary. If not otherwise stated, overnight cultures were diluted 1:20 into LB broth containing suitable antibiotics or additives like L-rhamnose and grown for 3 h at 37°C and 200 rpm before sample collection for downstream analyses.

### Generation of plasmids

Plasmids were generated by Gibson assembly^33^. For this purpose, vector fragments and inserts were amplified by PCR using the KAPA HIFI PCR Kit (Roche) and assembled using a Gibson Mix for 30 min at 50°C. The reaction product was transformed in *E. coli* Dh5α and selected on LB agar plates with appropriate antibiotics. Bacterial clones containing the correct plasmids were validated by sequencing (Eurofins). Primers for generation of plasmids are listed in **Table S4**.

### Generation of in-frame deletion and knock-in mutants

In-frame deletion mutants were generated using the suicide plasmid pEXG2 (67) as described in Klein et al.^34^ or its derivate pEXTK. In pEXTK, the *sacB* gene is replaced by a thymidine kinase gene. If pEXTK based mutator plasmids were used in the mutagenesis procedure, the positive selection to obtain a second crossover was performed by incubating merodiploidic clones for 3 h in 5 ml LB medium containing IPTG (1 mM). Subsequently, bacteria were positively selected by streaking bacteria on LB agar plates containing 200 μg/ml azidothymidine (Acros Organics) and 1 mM IPTG. Bacteria were then tested for loss of gentamicin sensitivity and mutants were verified by PCR as described previously^34^. The generated mutants and the primers used in this study are listed in **Table S3 or S4**, respectively.

### Generation of complementation constructs

Complementation of ID40 was done as described by Choi et al.^35^ The coding sequences of the gene of interest were amplified by PCR from genomic DNA and assembled with the plasmid pJM220 (pUC18T-miniTn7T-gm-rhaSR-PrhaBAD)^36^ by Gibson cloning. The constructed plasmids were transformed into *E. coli* SM10 λ pir and mobilized by conjugation into the mutant strains as described^35^ with some modifications. A triparental mating was conducted by combining the recipient strain together with the mini-Tn7T harbouring *E. coli* SM10 λ pir strain and *E. coli* SM10 λ pir pTNS3, harbouring a Tn7 transposase. Insertion of the mini-Tn7T construct into the *attTn7* site was verified by PCR. Excision of the pJM220 backbone containing the Gm resistance cassette was performed by expressing Flp recombinase from a conjugative plasmid, pFLP2. Finally, sucrose resistant but gentamicin and carbenicillin sensitive colonies were verified by PCR.

### RNA isolation and RT-qPCR

RNA isolation and RT-qPCR were performed as previously described ^34^. The used primers are listed in **Table S4**.

### Transcriptomics

The strains ID40 and ID40Δ*ygfB* were used to perform RNA sequencing and differential gene expression analysis (ID40 *vs* ID40Δ*ygfB*). The strains were subcultivated for 3 h in 5 ml LB medium. A total of four independent replicates per strain were used in the sequencing and analysis. RNA was isolated using the Quick-RNA™Fungal/Bacterial MiniprepKit (Zymo Research) according to manufacturer’s instruction. Subsequently, 15 μg RNA in 50 μl water was digested with 10 U DNAse I (Roche). The quality of the RNA was controlled by determination of the RNA Integrity Index using Agilent BioAnalyzer High Sensitivity DNA Assays. Successful depletion of DNA was controlled by qPCR and RT-qPCR for the *rpoS* and the *ygfB* genes. Next, the Zymo-Seq RiboFree total RNA Library Prep Kit (Zymo Research) was used to deplete ribosomal RNA and prepare samples for sequencing. For this step, 2 μg RNA per sample were used. Sequencing was performed with Illumina NextSeq500 (2×75 bp, MidOutput Flowcell). Mapping of sequencing reads and counting was performed using the subread package in R and the ID40 genome as a reference (https://www.ebi.ac.uk/ena/browser/view/LR700248)^37^. Differential gene expression analysis was performed using DeSeq2^38^.

### β-lactamase activity assay

A β-lactamase colorimetric activity assay (BioVision) based on nitrocefin turnover was performed according to manufacturers’ instructions after resuspending the bacteria in 5 μl/mg β-lactamase assay buffer and diluting the supernatant of sonicated bacteria 1:50 in β-lactamase assay buffer.

### AmpDh3 promoter-luciferase assays

To determine the activity of the *ampDh3*-promoter, various *ampDh3*-promoter-luciferase reporter constructs such as the plasmid pBBR-*ampDh3*-532-nanoluc were transformed into *Pseudomonas aeruginosa* strains via electroporation according to the protocol of Choi et al.^39^ (see **Table S3**).

Overnight cultures were subcultured for 3 h in 5 ml LB containing 75 μg/ml gentamicin. OD_600_ was measured and cultures were diluted to an OD_600_ of 0.2 in 1 ml LB. 50 μl were transferred into a white flat bottom 96 well plate in triplicates and 50 μl of Promega NanoGlo Luciferase assay reagent (Promega) prepared according to the manufacturer’s instructions was added to the wells. The plate was then shaken for 10 min at RT and chemiluminescence was measured using a Tecan Infinite Pro 200 plate reader.

### Determination of peptidoglycan catabolites in cytosolic fractions by LC-MS

*P. aeruginosa* ID40 parental and mutant strains were grown overnight in LB medium. The OD_600_ of overnight cultures (LB medium) was measured. 100 ml LB bacteria cultures with an initial OD_600_ of 0.05 per ml were grown for 6 h. Cells were harvested and OD_600_ measured. Bacteria were pelleted and resuspended in 20 ml 50 mM Tris-HCL buffer, pH 7.6 to a final concentration of OD_600_ = 5/ml (final OD_600_ =100). Subsequently, bacteria were centrifuged at 3,000 g for 10 min. The supernatant was discarded and the pellet frozen at -80°C. The next day, bacteria were resuspended in 400 μl water and the cultures boiled for 15 min at 95°C. Cultures were cooled down to and centrifuged at room temperature at 16,000 g for 10 min. 200 μl of the supernatant were added to 800 μl ice-cold acetone (MS grade, Sigma 34850-2.5L) to precipitate remaining proteins in the samples. Samples were then centrifuged at 4°C at 16,000 g for 10 min. The supernatant was transferred in a new tube and the cytosolic fraction was dried under vacuum for 2 h at 55°C in a Speedvac (Eppendorf). Pellets were then stored at 4°C. The dry cytosolic fractions were then dissolved in 50 μl Millipore water. 5 μl of the samples were subjected to LC-MS analysis with an UltiMate 3000 LC system (Dionex) coupled to an electrospray ionization-time of flight mass spectrometer (MicrO-TOF II; Bruker) that was operated in positive-ion mode in a mass range 180 m/z to 1,300 m/z. Metabolite separation was achieved with a Gemini C18 column (150 by 4.6 mm, 110 Å, 5 μm; Phenomenex) at 37°C with a flow rate of 0.2 ml/min in accordance with a previously described 45 min gradient program^40^ with small modifications: 5 min of washing with 100% buffer A (0.1% formic acid, 0.05% ammonium formate in water), followed by a linear gradient over 30 min to 40% buffer B (acetonitrile) and a 10 min column re-equilibration step with 100% buffer A. Peptidoglycan (PG) metabolites were shown in Data Analysis (Bruker) by extracted ion chromatograms (EICs) and the area under the curves of the respective EICs were calculated in Prism 8 (GraphPad). The theoretical m/z values of the PG metabolites investigated are 276.108 m/z for anhMurNAc, 479.187 m/z for GlcNAc-anhMurNAc, 648.272 m/z for anhMurNAc-3P, 851.352 m/z for GlcNAc-anhMurNAc-3P, 790.347 m/z for anhMurNAc-5P, 680.110 m/z for UDP-MurNAc, and 1194.349 (597.678 ^2+^) for UDP-MurNAc 5P.

### Induction of DNA damage with ciprofloxacin

Induction of DNA-damage using ciprofloxacin was adapted from Peña et al.^16^. Overnight cultures were subcultured in LB for 3 h at 37°C and grown until exponential phase. The cultures were diluted to OD_600_ 0.5 and 32 μg/ml ciprofloxacin (Sigma-Aldrich) was added to the cultures if not otherwise stated. Cultures were incubated for two hours at 37°C and harvested by centrifuging appropriate cell numbers for the desired downstream analyses.

### Antibiotic susceptibility testing

For antibiotic susceptibility testing by microbroth dilution, bacterial strains were grown overnight at 37°C in LB medium. Physiological NaCl solution was inoculated to a McFarland standard of 0.5. Subsequently 62.5 μl of the suspension were transferred into 15 ml MH broth and mixed well. According to the manufacturer’s instructions, 50 μl of the suspension was transferred into each well of a microbroth dilution microtiter plate (Sensititre™ GN2F, Sensititre™ EUX2NF (Thermo Fisher Scientific)). Microtiter plates were incubated for 18 h at 37°C and OD_600_ was measured using the Tecan Infinite® 200 PRO. Bacterial strains were considered as sensitive to the respective antibiotic concentration if an OD_600_ value below 0.05 was measured.

### Checkerboard assay

Stocks of antibiotics to test were prepared by dissolving them according to CLSI M100 Performance Standards for Antimicrobial Susceptibility Testing^41^ in the indicated solvent and diluent to a final concentration of 5.12 mg/ml. Salts were corrected for their mass. Used antibiotics: Ciprofloxacin hydrochloride monohydrate (Sigma-Aldrich; European Pharmacopoeia Reference Standard), piperacillin sodium (Sigma-Aldrich, analytical standard), imipenem (Sigma-Aldrich; European Pharmacopoeia Reference Standard), ceftazidime pentahydrate (Sigma-Aldrich; European Pharmacopoeia Reference Standard) and aztreonam (United States Pharmacopeia Reference Standard).

Working stocks were then prepared by serial dilution in MHB II medium. Plates for checkerboards were prepared by adding 25 μl of each antibiotic at 4x the final concentration to be tested in the respective well in a flat bottom, transparent 96 well plate (Greiner). In one column a growth control was prepared by adding 50 μl of MHB II medium. A sterility control was prepared in a second column by adding 100 μl of MHB II medium. Inocula of the strain to be tested were prepared by inoculating physiological NaCl solution to a McFarland standard of 0.5 from overnight cultures. 125 μl of this solution were then added to 15 mL of MHB II medium and 50 μl of this inoculum added to the wells containing antibiotics as well as to the growth control wells. Plates were incubated for 20 hours at 37°C. After incubation the OD_600_ values were determined using a Tecan Infinite® 200 PRO. Each assay was prepared in duplicate. For each replicate, the ratio of signal for each well and the mean of the sterility control was calculated. The mean value of both replicates was calculated. If the value was smaller than 1.5, this concentration was considered to be inhibitive. From these values, the MIC and FIC-I for the tested antibiotics were calculated as follows: FIC_A_ = MIC_A combined_ / MIC_A single antibiotic_; FIC_B_ = MIC_B combined_ / MIC_B single antibiotic_; FIC-I = FIC_A_ + FIC_B_.

Interpretation of FIC-I: ≤0.5: Synergism; >0.5 to 1: additive effect; >1 to 4: indifferent effect; >4: Antagonism

### Expression and purification of His-MBP-AlpA and His-MBP

For purification of His-MBP-AlpA and His-MBP, expression cultures of 1 liter LB medium were inoculated at an OD_600_ of 0.15 with starter cultures of *E. coli* BL21 carrying either pETM-41_AlpA or pETM-41_stop. Expression cultures were grown until an OD_600_ of 0.6-0.8 at 37°C. The cultures were then shifted to 20°C and equilibrated for 30 min. IPTG was added to a final concentration of 1 mM and expression was carried out at 20°C for 18 h. Cultures were harvested by centrifuging at 6,000 g for 10 min at 4°C.

Pellets were resuspended in 35 ml lysis buffer (50 mM Tris, 150 mM NaCl, 25 mM imidazole, pH 7.5) supplemented with lysozyme, Triton X-100, DNase and cOmplete protease inhibitor cocktail (Roche). Bacteria were lysed by sonication for 3 x 1 min on ice at 20% amplitude and 50% duty cycle. Cell debris was removed by centrifuging the lysate at 35,000 g for 1 h at 4°C. The supernatant was sterile filtered through a 0.22 μm syringe filter (Millipore) and affinity-purified in a gravity flow column using Ni^2+^-NTA-agarose beads (Qiagen). After binding of the His-tagged proteins to the columns, columns were washed with lysis buffer and proteins were eluted using elution buffer (lysis buffer + 350 mM imidazole). Fractions were analyzed via SDS-PAGE and Coomassie staining. Elution fractions were dialyzed in 3 liter dialysis buffer (50 mM Tris, 150 mM NaCl, 20% V/V glycerol at pH 7.5) using Slide-A-Lyzer dialysis cassettes (ThermoFisher) with 20 kDa cutoff and 12-30 ml volume. Pure protein was aliquoted and stored at -80°C after snap freezing with liquid nitrogen.

### Expression and purification of GST, GST-YgfB

Expression and purification were performed as above, using the strain *E. coli* BL21 carrying either carrying pGEX4T3_stop or pGEX4T3_YgfB. Differing from above, the expression was carried out at 25°C. For resuspension and lysis of the bacterial pellet, GST-A buffer (50 mM Tris, 150 mM NaCl, 1 mM DTT, pH 7.5) supplemented with lysozyme, Triton X-100, DNase and protease inhibitor was used. For purification, a GSTrap™ HP 1 ml column (Cytiva) connected to a peristaltic pump was used. After loading the column and collecting the flow through the column was washed using GST-A buffer and the protein eluted using GST-B-buffer (50 mM Tris, 150 mM NaCl, 10 mM reduced glutathione, pH 8). After column regeneration, the flow through was loaded on the column once again and also washed and eluted. The obtained eluate fractions were pooled and dialysed against 10 liter of PBS pH 7.4 and 0.5 mM DTT using a ZelluTrans (Roth) dialysis tube with a 3.4 kDa cutoff and frozen in dialysis buffer. Analysis by SDS-PAGE and protein storage was done as described above.

### Expression and purification of YgfB

Expression and purification were performed similar to as described for His-MBP and His-MBP-AlpA, using the strain *E. coli* BL21 carrying pETM-30_YgfB, however, the expression was carried out at 25°C. This purification step yielded His-GST-TEV-YgfB. Then, His-tagged TEV-protease was added to the elution fraction containing His-GST-TEV-YgfB and dialyzed in 2 l dialysis buffer (50 mM Tris, 150 mM NaCl, 1 mM DTT at pH 7.5) using a ZelluTrans (Roth) dialysis tube with a 3.4 kDa cutoff over night at 4°C.

The yielded cleavage product was purified using reverse Ni^2+^-affinity chromatography using Ni^2+^-NTA-agarose beads equilibrated with sterile filtered dialysis buffer and the flow through containing only YgfB was collected, aliquoted and stored as above. Fractions were analyzed via SDS-PAGE and Coomassie staining.

### Generation of affinity purified antibodies against recombinant GST-YgfB

Antibodies were raised in 2 rabbits using recombinant GST-YgfB. Those obtained from one rabbit were selected for best performance against ID40, ID40Δ*ygfB* and recombinant GST-YgfB, and subsequently affinity-purified against GST-YgfB protein (Eurogentec).

### Quantification of HiBiT-tagged proteins using a luciferase assay

Subcultures were harvested by centrifuging at 5,000 g for 10 min. Cell pellets were washed once by resuspending in 1 ml PBS and centrifuged at 10,000 g for 1 min. The pellet was resuspended 1 ml PBS and the OD_600_ was measured. Bacteria corresponding to an OD_600_ = 1 were harvested by centrifugation at 10,000 g for 1 min. The pelleted bacteria were resuspended in 500 μl buffer K adapted from Dietsche et al.^42^ (50 mM triethanolamine pH 7.5, 250 mM sucrose, 1 mM EDTA, 1 mM MgCl2, 0.5% Triton-X 100, 10 μg/ml DNase, 20 μg/ml lysozyme, 1:100 cOmplete protease inhibitor cocktail) and incubated on ice for 30 min. For quantification of HiBiT-tagged proteins, 50 μl Nano-Glo HiBiT Lytic Reagent containing 1 μl furimazine and 2 μl recombinant LgBiT were added to 50 μl lysate in a white flat-bottom 96 well plate (Greiner) in technical triplicates. Plates were incubated for 10 min and chemiluminescence was measured using a Tecan Reader Infinite 200 Pro plate reader (500 ms integration time).

### Western blot analyses from whole cell lysates

After treating bacteria according as desired, an equivalent of OD_600_ = 10 of bacteria was boiled in 2x Laemmli Sample Buffer (BioRad #1610747) supplemented with 5% 2-mercaptoethanol for 10 min at 95°C. Of the whole cell lysates, 10 μl were loaded onto a Mini Protean TGX Precast Protein gel (BioRad) with the acrylamide percentage chosen according to the protein to be detected. After separation of the proteins, they were transferred onto a nitrocellulose membrane (Amersham). HiBiT-tagged proteins were detected using the Nano-Glo HiBiT blotting system (Promega) according to the manual, while other proteins were detected via antibodies.

For this purpose, the membrane was blocked with 1x BlueBlock PF (Serva) and afterwards incubated with the primary antibody in 1x BlueBlock solution (rabbit anti-YgfB, 1:500; rabbit anti-HA-Tag, 1:1,000 (CellSignaling; HA-Tag (C29F4) Rabbit mAb #3724); mouse anti-RpoB (*Ec*), 1:1,000 (BioLegend; Anti-*E. coli* RNA Polymerase β Antibody Mouse); Rabbit anti-SurA, 1:200^34^). Next, membranes were washed three times with TBS-T (50 mM TRIS, 150 mM NaCl pH 7.4 with added 0.1% Tween-20) and afterwards incubated with the appropriate secondary antibodies (horseradish-peroxidase-conjugated goat anti-rabbit antibody 1:2,000 (Dianova; Goat F(ab’)2 anti-Rabbit IgG (H+L)-HRPO, MinX none) or horse-radish-peroxidase-conjugated anti-mouse antibody, 1:2,000 (Invitrogen; Rabbit anti-Mouse IgG (H+L) Secondary Antibody, HRP)). The membranes were washed again three times using TBS-T. For detection, Clarity Western ECL Substrate (BioRad) was added to the membrane and the signal was detected by a Fusion Solo S imager (Vilber). The anti-RpoB signal was used as loading control.

### Subcellular fractionation of *Pseudomonas aeruginosa* by spheroblasting

For subcellular fractionation, a modified version of the protocol described by Wang et al.^43^ was used. *P. aeruginosa* strains were grown in LB medium overnight. The cultures were inoculated into fresh LB with the OD_600_ adjusted to 0.05 and subcultivated at 37°C with shaking to an OD_600_ = 1.5. 10 ml of each strain were harvested by centrifugation at 4,500 g for 10 min. The cell pellets were resuspended in 500 μl sucrose-EDTA solution (2.5 mM EDTA and 20% (w/v) sucrose in PBS, pH 7.3) and incubated at room temperature for 20 min. 500 μl ice-cold H_2_O were added and the samples were incubated for 5 min at 4°C with gentle shaking (550 rpm). After centrifugation for 20 min at 4°C and 7,000 g, the supernatant containing all periplasmic proteins was removed and filtered through a syringe filter with 0.2 μm pore size. To obtain the cytosolic and membrane-bound proteins, the pellets were resuspended in 375 μl H_2_O and 125 μl 4x Laemmli solution with 10% β-mercaptoethanol and then incubated for 10 min at 95°C.

To obtain the periplasmic proteins, 250 μl of 14.3% aqueous trichloroacetic acid solution (w/v in H_2_O) were added to 1 ml of the supernatants containing the periplasmic proteins and incubated on ice for 30 min. After centrifugation for 5 min at 4°C and 14,000 g, the supernatant was discarded. The pellets were washed twice by adding 400 μl acetone, centrifuging at 14,000 g and 4°C for 5 min and discarding the acetone. The pellets were dried at 95°C for 1 min, then resuspended in 36 μl H_2_O. 12 μl 4x Laemmli buffer with 10% β-mercaptoethanol were added and the samples incubated for 10 min at 95°C prior to loading on an SDS polyacrylamide gel.

### GST pull-downs from cell lysates

Day cultures were inoculated at an OD_600_ of 0.1 in 500 ml of LB with overnight cultures of ID40Δ*ygfB*::*alpA*-HiBiT::HA-*alpR* and grown for 5 h at 37°C. Cultures were harvested by centrifuging for 10 min at 6,000 g. Cell pellets were resuspended in 5 ml pulldown-buffer (50 mM Tris pH 7.5, 300 mM NaCl, 0.5% IGEPAL, 2 mM DTT) supplemented with cOmplete protease inhibitor cocktail, DNase I, lysozyme and Triton-X 100. Cells were lysed by sonification, cell debris removed by centrifugation and supernatants were used for downstream application after sterile filtration.

1 ml of recombinant GST or GST-YgfB protein at a concentration of 10 μM was incubated with 100 μl 50 % MagneGST (Promega) bead-slurry equilibrated with pulldown-buffer for 45 min at 4°C and washed two times with 500 μl pulldown-buffer. The beads were then incubated with 1 ml of cell lysate for 45 min at 4°C. After washing the beads three times with 700 μl pulldown-buffer, the bound proteins were eluted from the beads using 100 μl pulldown-buffer supplemented with 25 mM glutathione. 33 μl of 4x Laemmli buffer was added to the eluate and the samples were boiled for 10 min at 95°C. For the input samples, 10 μl of GST-tagged protein was mixed with 10 μl of cell lysate. 20 μl 4x Laemmli buffer was added and samples were boiled at 95°C for 10 minutes. Samples were analysed by SDS-PAGE and Western Blot using the Nano-Glo HiBiT blotting system (Promega).

### Pulldowns using His-tagged recombinant proteins

1 ml of recombinant His-MBP-AlpA and His-MBP at a concentration of 10 μM was incubated with 100 μl MagneHis™ Ni Particles (Promega) equilibrated with pulldown-buffer (50 mM Tris pH 7.5, 25 mM imidazole, 300 mM NaCl, 0.5% IGEPAL, 2 mM DTT) for 45 min at 4°C and washed 2 times with 500 μl pulldown-buffer. 1 ml of recombinant YgfB at a concentration of 10 μM was added to the beads and incubated for 45 min at 4°C. After washing the beads three times with 700 μl pulldown-buffer, the bound proteins were eluted with 75 μl pulldown-buffer supplemented with 350 mM imidazole. 25 μl 4x Laemmli buffer were added to the eluate and the samples were boiled for 10 min at 95°C. For the input samples, 10 μl of His-tagged protein was mixed with 10 μl of rYgfB. After addition of 20 μl 4x Laemmli buffer the samples were boiled at 95°C for 10 min. Proteins were detected by SDS-PAGE and Western blot as described above.

### Electromobility Shift Assays (EMSA)

For generation of the labeled probe, 5’-IRDye® 700 labeled oligonucleotides were purchased from IDT with the following sequences: ABE; 5’-CGG TGT TGC ACG CGG C**GG GAC GCT CGC GGT AGT TT**T TTC CCA TGA TCA CG-3’ and 5’-CGT GAT CAT GGG AAA **AAA CTA CCG CGA GCG TCC C**GC CGC GTG CAA CAC CG-3’ and scrambled control probes; 5’-GTT TAC TAG GTC GAG GTA CTT CGA CGC GCG CCG TCT GCT AGC GCG GTC TG-3’ and 5’-CA GAC CGC GCT AGC AGA CGG CGC GCG TCG AAG TAC CTC GAC CTA GTA AAC-3’ The AlpA binding element is shown in bold letters. The oligonucleotides were annealed by mixing them in equimolar amounts in duplexing buffer (100 mM Potassium Acetate; 30 mM HEPES, pH 7.5) and heating to 100°C for 5 min in a PCR cycler. The cycler was then turned off and the samples were allowed to cool to room temperature while still inside the block. The annealed product was then diluted with water to 6.25 nM for EMSA experiments.

For EMSAs fluorophore labeled DNA probes at a concentration of 0.3125 nM were incubated for 30 min at 20°C in 20 μl reaction mix (Licor Odysee EMSA Kit) containing 33.4 mM Tris, 70.2 mM NaCl, 12.5 mM NaOAc, 3.75 mM HEPES, 50 mM KCl, 3.5 mM DTT, 0.25% Tween-20 and 0.5 μg sheared salmon sperm DNA (ThermoFisher) with proteins. For resolving the reactions, 4% polyacrylamide gels containing 30% triethylene glycol were cast (For two gels: 2 ml ROTIPHORESE®Gel 30 37.5:1 (Roth), 4.5 ml triethylene glycol (Sigma-Aldrich), 1.5 ml 5x TBE-buffer, 7 ml ddH_2_O, 15 μl TEMED, 75 μl 10% APS). The gels were preequilibrated for 30 minutes at 130 V in 0.5x TBE-buffer. Samples with added 10x orange dye were then loaded onto the gel at 4°C and the voltage set to 300 V until the samples entered the gel completely. The voltage was then turned down to 130 V and the gel was run until the migration front reached the end of the gel. The gels were imaged using the Licor Odyssey imaging system using the 700 nm channel.

### Statistics and reproducibility

Statistics were performed using GraphPad Prism 9.12 software as described for each experiment in the table or figure legends. Depicted are mean and standard deviation of *n* replicates as indicated in the figure legend. Significance levels were denoted as: ns p>0.05, *p<0.05, **p<0.01, ***p<0.001 ****p<0.001. Data distribution was checked for normal distribution using Shapiro-Wilks test. Promoter luciferase assay data and LC-MS data were transformed to log_10_ since they were lognormal distributed.

## Data availability

Transcriptomic data is available on the server of the National Library of Medicine under the accession number PRJNA835697 Reviewer link: https://dataview.ncbi.nlm.nih.gov/object/PRJNA835697?reviewer=fe3gcngqnecij22uk4lm999mte. Genomic DNA sequence of ID40 is available under https://www.ebi.ac.uk/ena/browser/view/LR700248. Primary data are available from the corresponding author upon request.

## Acknowledgments

This work was financed by Grants from the Deutsche Forschungsgemeinschaft (DFG) (BO 1527/4-1) to EB/MS; the German Center of Infection Research (DZIF) to MS and the DFG Cluster of Excellence EXC2124 ‘Controlling Microbes to Fight Infection’ (CMFI) to CM/EB. LM was supported by the intramural IZKF program of the Medical Faculty at the University of Tübingen. We thank Libera Lo Presti for the critical reading of the manuscript, fruitful comments and great corrections.

## Contribution of authors

Study design EB, MS, FR, OE, CM, MB; manuscript and figures by EB, MS, FR, OE, MB, VE, CM; transcriptomics AA, CE, EW, EB; analysis of PG precursors by MB, EB, CM; generation of mutants by EB, EW, FS, KK, NW, OE, LM, VE, MS; protein purification and antibody generation by FR, OE, FS, KK; protein-protein interaction by OE, FS; EMSAs by OE, Western blots by OE, FS, NW; localization of AmpDh3 by VE; *ampDh3* promoter analysis by EB, EW, LM, NW; Quantification of HiBiT tagged proteins by EB, OE; in silico analysis FR; antibiotic resistance and β-lactamase activity by EW, NW, OE, LM, MS; checkerboard assays: OE, FS

