## Supplementary information for "YgfB increases β-lactam resistance in *Pseudomonas aeruginosa* by counteracting AlpA-mediated *ampDh3* expression"

### Supporting information

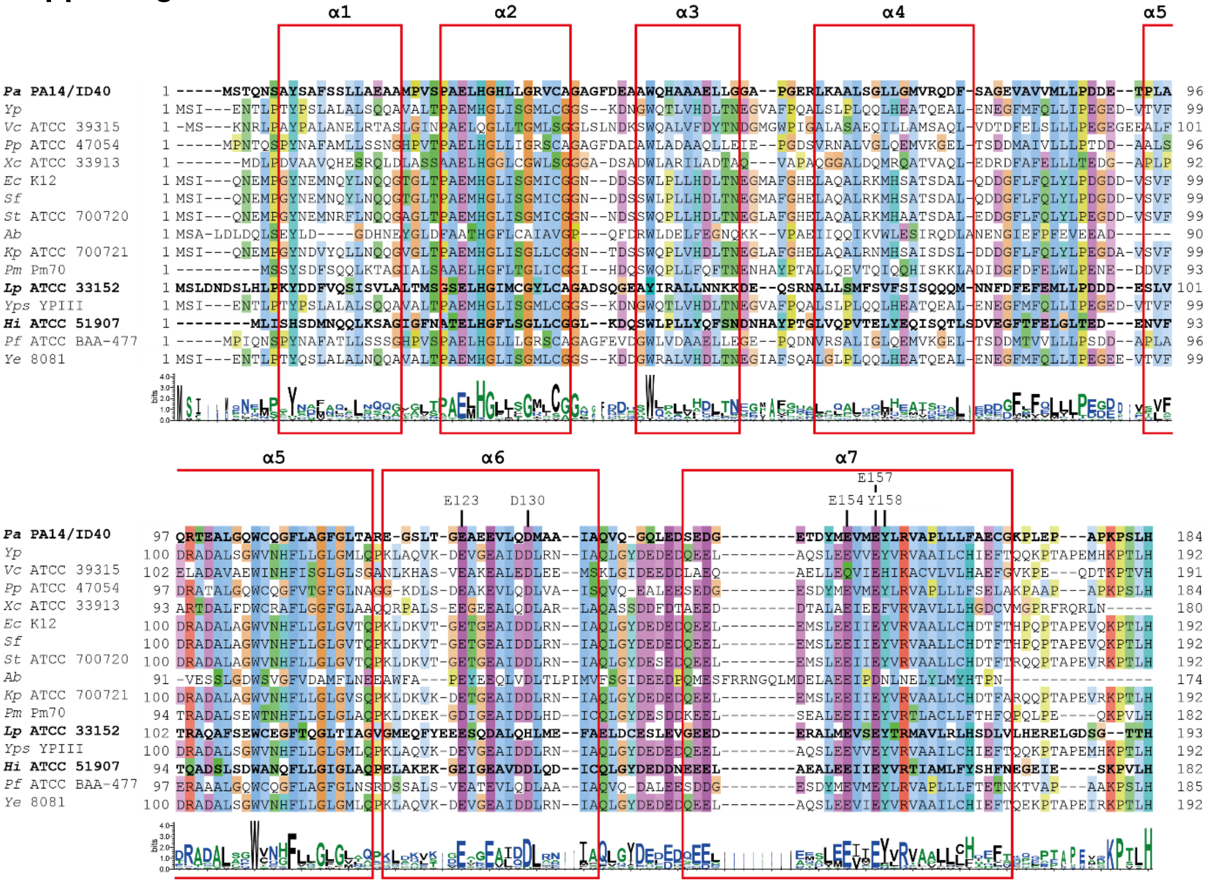

**Fig. S1: Conservation of YgfB in different species.** Sequences of YgfB proteins were aligned using clustalΩ[1, 2] and color coded according to conservation with clustalX[3]. The secondary structure elements as deduced from the crystal structure of the *Haemophilus influenzae (Hi)* protein (pdb ID: 1izm) are indicated as red boxes. Conserved residues in the dimerization interface of *Hi* YgfB[4] are labeled above the alignment. The sequences of the *Pseudomonas aeruginosa* (*Pa*) protein as well as the sequences of the crystal structures of the *Legionella pneumophila* (*Lp*, pdb ID: 4gyt) and *Hi* proteins are highlighted in bold. A sequence logo generated with WebLogo 3[5, 6] is shown below the sequence alignment to highlight conserved residues. The sequence alignment view was prepared with Jalview[7] UniProt accession numbers and species names are listed in **Table S5**.

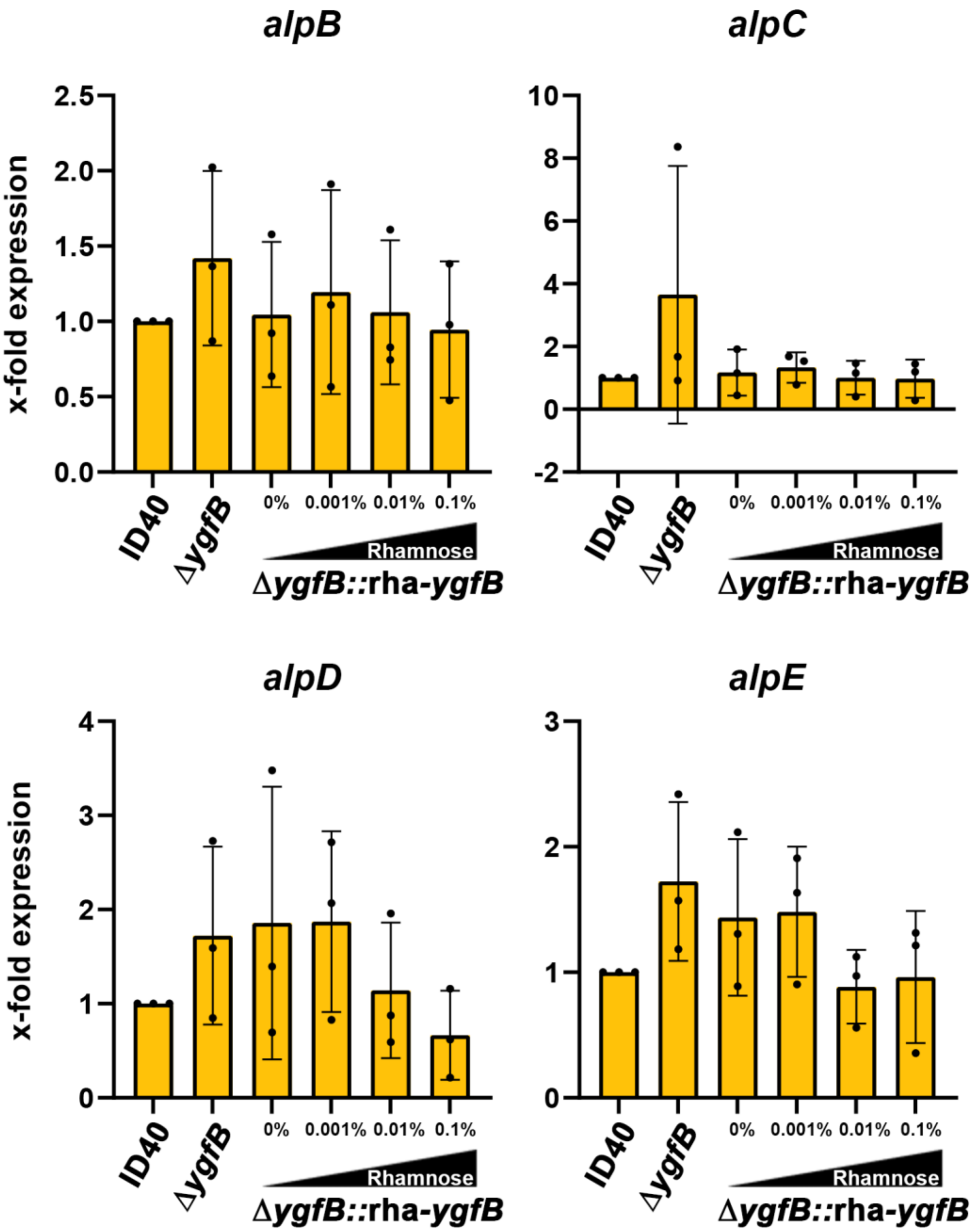

**Fig. S2: Gene expression of the alp*BCDE* cluster.** ID40, ID40Δ*ygfB,* ID40Δ*ygfB*::rha-*ygfB* were grown with the indicated concentrations of rhamnose in LB medium at 37°C to induce *ygfB* expression. After 3h of growth mRNA was isolated and used for RT-qPCR. Data depict mean and SD for x-fold expression of the indicated genes compared to ID40 of *n*=3 independent experiments.

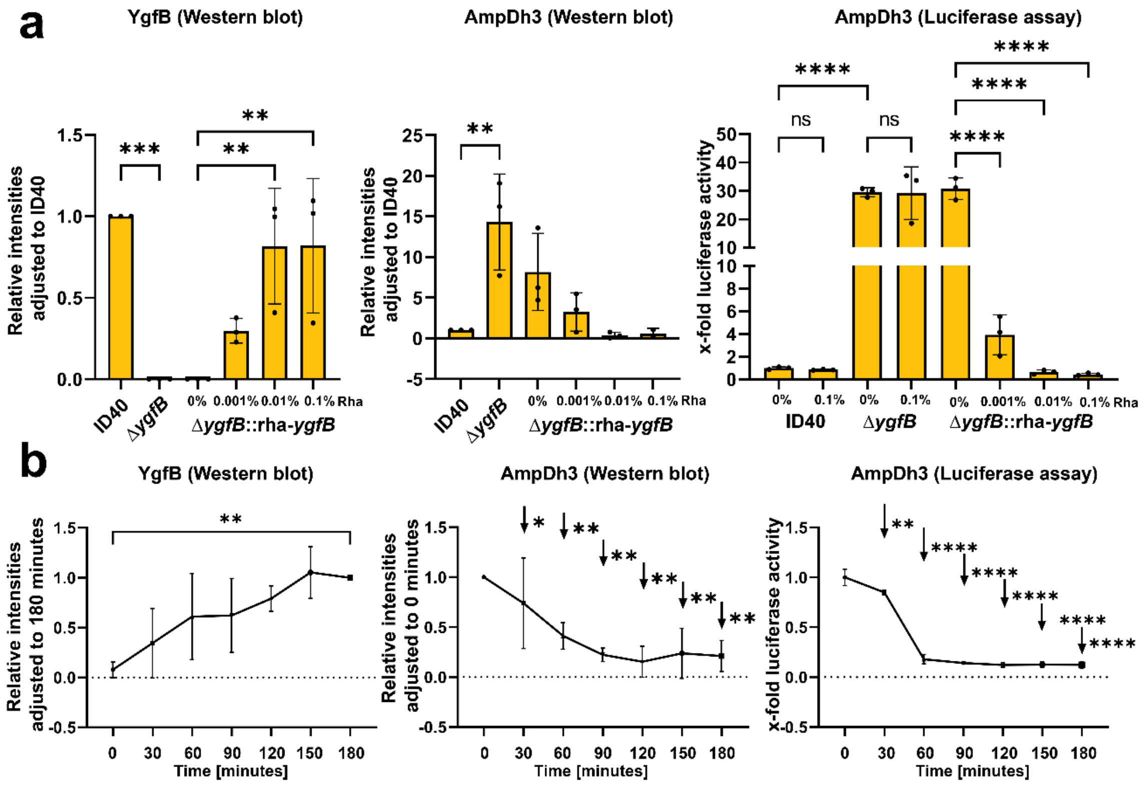

**Fig. S3: Semiquantification of YgfB and AmpDh3 based on WB analyses and of AmpDh3-HiBiT by luciferase assay.**

Western blots shown in (a) Fig 1b and (b) Fig. 2b were quantified using Image J. Bands for YgfB and AmpDh3 were normalized to RpoB (left and middle panels), *n*=3. In the right panel experiments are shown which were performed with the same strains and under the same conditions as the WB analyses but the cells were lysed with a lysis buffer. The lysates were incubated with LgBiT and furimazine to measure relative AmpDh3-HiBiT levels as luciferase activity in a Tecan reader (*n*=3). Of note: in (a), right panel, ID40 and Δ*ygfB* strains were in addition treated with 0.1% rhamnose to demonstrate that rhamnose has no impact on AmpDh3 levels.

Statistical analyses were performed for WT vs. Δ*ygfB* with or without addition of 0.1% rhamnose and the *ygfB* complemented strain without vs. with rhamnose as well as time point zero vs. all other time points. (Asterisks indicate significant differences, ns: not significant, *p<0.05, **p<0.01,***p<0.001, ****p<0.0001, in (a), left panel, comparing to timepoint 180 minutes and in (a), middle panel, comparing to timepoint 0 minutes, one-way ANOVA, Šídák's multiple comparisons).

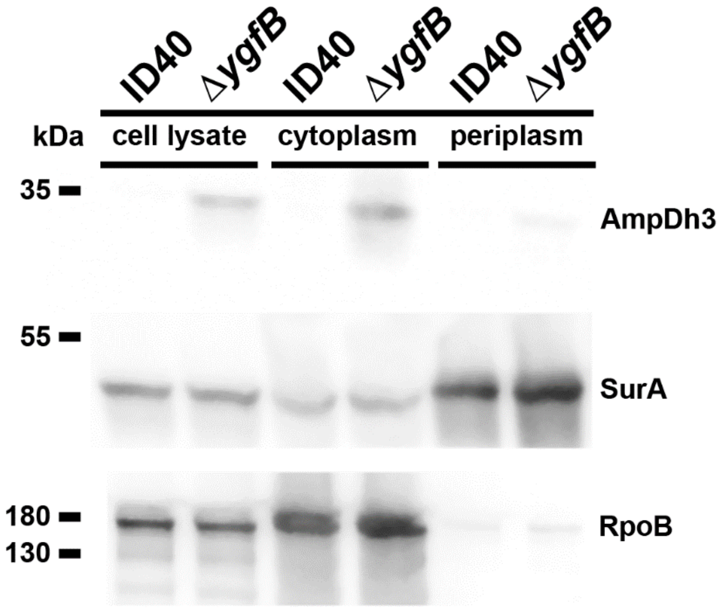

**Fig. S4: Localization of AmpDh3.** Subcellular fractionation using the strains ID40:*:ampDh3*-HiBiT and ID40Δ*ygfB::ampDh3*-HiBiT was performed as described in material and methods. Western blots were performed for AmpDh3-HiBiT, SurA and RpoB. Left panel: total cells boiled in Laemmli buffer (input), Middle Panel: cytoplasmic fraction; Right panel: periplasmic fraction.

**Table S1. Fractional inhibitory concentration index (FIC-I).** FIC-I ≤ 0.5, synergistic effect, 0.5-1 additive, >1 indifferent.

| | CAZ + CIP | | | PIP + CIP | | |
|---|---|---|---|---|---|---|
| **Strain** | **$FIC_{CAZ}$** | **$FIC_{CIP}$** | **FIC-I** | **$FIC_{PIP}$** | **$FIC_{CIP}$** | **FIC-I** |
| ID40 | 0.25 | 0.25 | 0.5 | 0.25 | 0.5 | 0.75 |
| Δ*ygfB* | 0.25 | 0.25 | 0.5 | 0.25 | 0.25 | 0.5 |
| Δ*ampDh3* | 0.25 | 0.25 | 0.5 | 0.25 | 0.5 | 0.75 |
| Δ*ygfB*Δ*ampDh3* | 0.25 | 0.25 | 0.5 | 0.25 | 0.5 | 0.75 |
| Δ*alpA* | 0.125 | 0.25 | 0.375 | 0.0625 | 0.25 | 0.3125 |
| Δ*ygfB*Δ*alpA* | 0.125 | 0.25 | 0.375 | 0.125 | 0.25 | 0.375 |
| Δ*ygfB*::rha-*ygfB* 0% rha | 0.25 | 0.25 | 0.5 | 0.125 | 0.5 | 0.625 |
| Δ*ygfB*::rha-*ygfB* 0.1% rha | 0.25 | 0.25 | 0.5 | 0.25 | 0.5 | 0.75 |
| Δ*ampDh3*::rha-*ampDh3* 0% rha | 0.125 | 0.5 | 0.625 | 0.25 | 0.5 | 0.75 |
| Δ*ampDh3*::rha-ampDh3 0.1% rha | 0.015625 | 0.5 | 0.515625 | 0.5 | 0.5 | 1 |
| Δ*alpA*::rha-*alpA* 0%rha | 0.25 | 0.25 | 0.5 | 0.125 | 0.5 | 0.625 |
| Δ*alpA*::rha-*alpA* 0.1%rha | 0.25 | 0.25 | 0.5 | 0.25 | 0.125 | 0.375 |

| | IMP + CIP | | | AZT + CIP | | |
|---|---|---|---|---|---|---|
| **Strain** | **$FIC_{IMP}$** | **$FIC_{CIP}$** | **FIC-I** | **$FIC_{AZT}$** | **$FIC_{CIP}$** | **FIC-I** |
| ID40 | 0.25 | 0.5 | 0.75 | 0.25 | 0.25 | 0.5 |
| Δ*ygfB* | 0.0625 | 0.5 | 0.5625 | 0.25 | 0.25 | 0.5 |
| Δ*ampDh3* | 0.5 | 0.25 | 0.75 | 0.25 | 0.25 | 0.5 |
| Δ*ygfB*Δ*ampDh3* | 0.25 | 0.5 | 0.75 | 0.03125 | 0.5 | 0.53125 |
| Δ*alpA* | 0.125 | 0.5 | 0.625 | 0.25 | 0.25 | 0.5 |
| Δ*ygfB*Δ*alpA* | 0.125 | 0.5 | 0.625 | 0.125 | 0.125 | 0.25 |
| Δ*ygfB*::rha-*ygfB* 0% rha | 0.125 | 0.5 | 0.625 | 0.25 | 0.25 | 0.5 |
| Δ*ygfB*::rha-*ygfB* 0.1% rha | 0.25 | 0.5 | 0.75 | 0.5 | 0.25 | 0.75 |
| Δ*ampDh3*::rha-*ampDh3* 0% rha | 0.25 | 0.5 | 0.75 | 0.5 | 0.25 | 0.75 |
| Δ*ampDh3*::rha-ampDh3 0.1% rha | 0.03125 | 0.5 | 0.53125 | 0.5 | 0.25 | 0.75 |
| Δ*alpA*::rha-*alpA* 0%rha | 0.25 | 0.5 | 0.75 | 0.25 | 0.25 | 0.5 |
| Δ*alpA*::rha-*alpA* 0.1%rha | 0.03125 | 0.5 | 0.53125 | 0.5 | 0.25 | 0.75 |

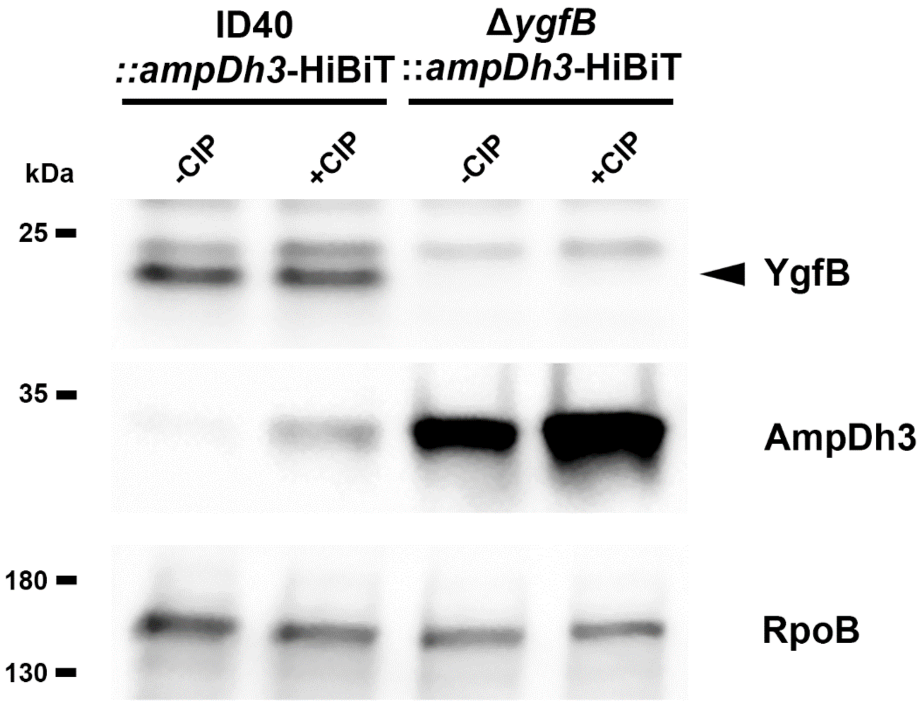

**Fig. S5: Production of AmpDh3 upon stimulation with subinhibitory concentrations of ciprofloxacin (CIP).** The indicated strains were adjusted to a McFarland of 0.5 in LB medium and then treated with or without 2.5 µg/ml CIP for 18 h. Whole cells were lysed in Laemmli buffer. Protein detection was performed either using recombinant LgBiT and furimazine for AmpDh3-HiBiT or anti-RpoB and anti-YgfB, respectively, followed by HRP-conjugated secondary antibody and ECL as substrate. Blots shown are representative for three independent experiments.

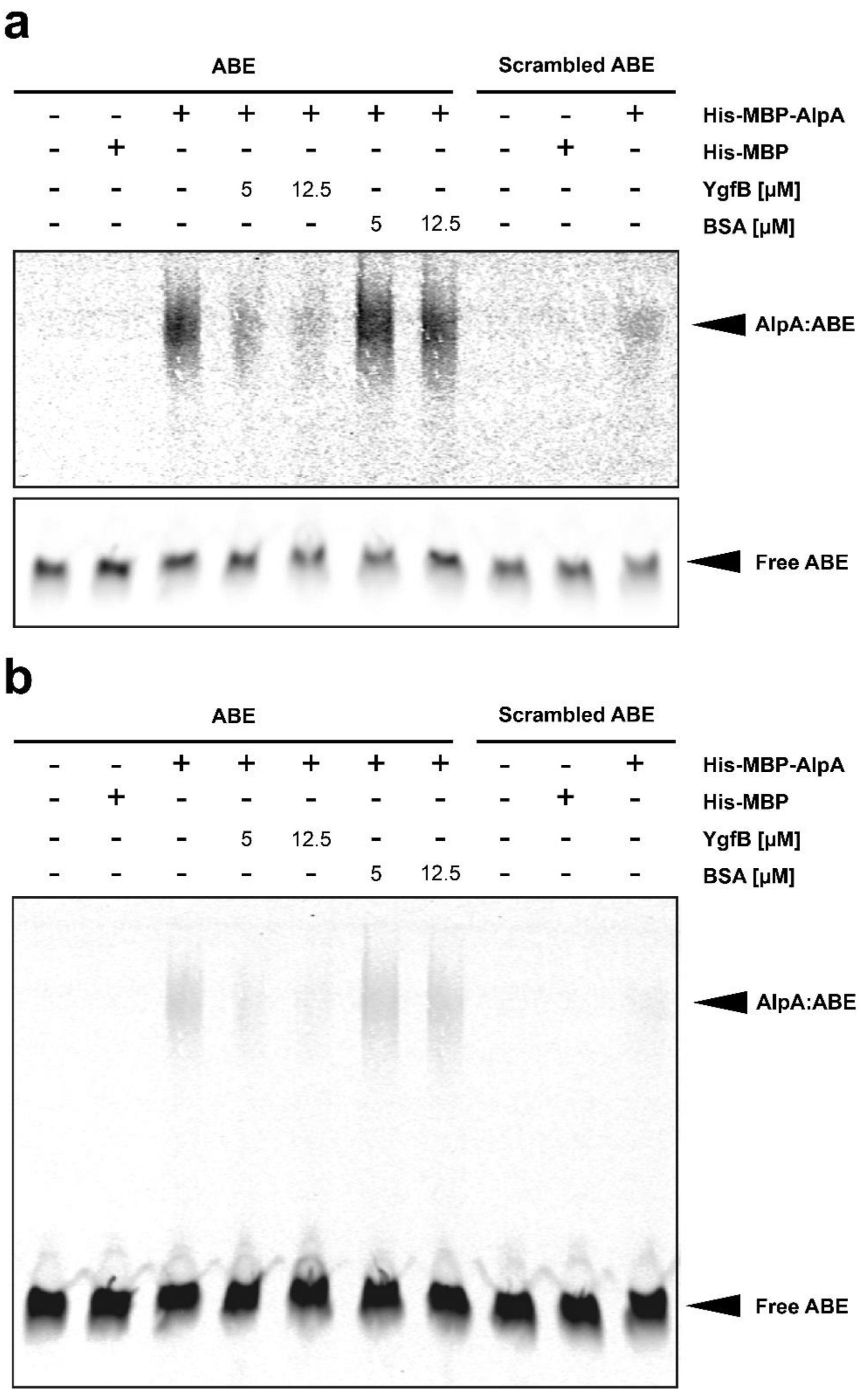

**Fig. S6: Interaction of AlpA with the AlpA binding element (ABE) of the *ampDh3* promoter.** (a) EMSA using 0.3125 nM IRDye700-labeled AlpA binding element (ABE) or 0.3125 nM IRDye700-labeled scrambled DNA probe incubated with the indicated proteins. The assay shown here is representative of five experiments. Concentrations used: His-MBP-AlpA and His-MBP 1.25 µM; YgfB and BSA 5 µM and 12.5 µM. Detection of IRDye700 was performed using the Licor Odyssey detection system. Of note, to obtain a higher dynamic range the free dsDNA and the shifted DNA-protein complexes were recorded separately. (b) In this case fluorescence was recorded over the entire gel to simultaneously record free ABE and AlpA:ABE complex.

**Table S2: MIC values for the indicated strains (in mg/l).** MEM meropenem, IMP imipenem, FEP cefepime, CAZ ceftazidime, PIP piperacillin, TZP tazobactam/piperacillin, AZT aztreonam, CIP ciprofloxacin. Green: reduced MIC compared to WT, Green bold and green coloured field: resistance is broken, nd not determined.

| | MEM | IMP | FEP | CAZ | PIP | TZP | AZT | CIP |
|---|---|---|---|---|---|---|---|---|
| MIC S< | **2** | **4** | **8** | **8** | **16** | **16** | **16** | **0.5** |
| MIC R> | **8** | **4** | **8** | **8** | **16** | **16** | **16** | **0.5** |
| **ID143 WT** | 32 | 32 | 16 | 32 | >128 | >128 | >32 | >4 |
| **ID143Δ*ygfB*** | 2-4 | **2** | **8** | **<2-2** | **<16** | **8** | **4** | >4 |
| **ID72 WT** | 16 | 32 | 32 | >32 | >128 | >128 | 32 | <0.125 |
| **ID72Δ*ygfB*** | **2** | **4** | **2** | **4** | **<16** | **<8** | **8** | 0.125 |
| **PAO1** | **<0.5** | 8 | **1** | **<1-1** | **<16** | **<4** | **2** | <0.125 |
| **PAO1Δ*ygfB*** | **<0.5** | **<1** | **1** | **<1-1** | **<16** | **<4** | **1** | <0.125 |
| **PA14 WT** | **<0.5** | **<1** | **<1** | **2** | **<4** | nd | **4-8** | <0.125 |
| **PA14Δ*ygfB*** | **<0.5** | **<1** | **<1** | **2** | **<4** | nd | **4** | <0.125 |

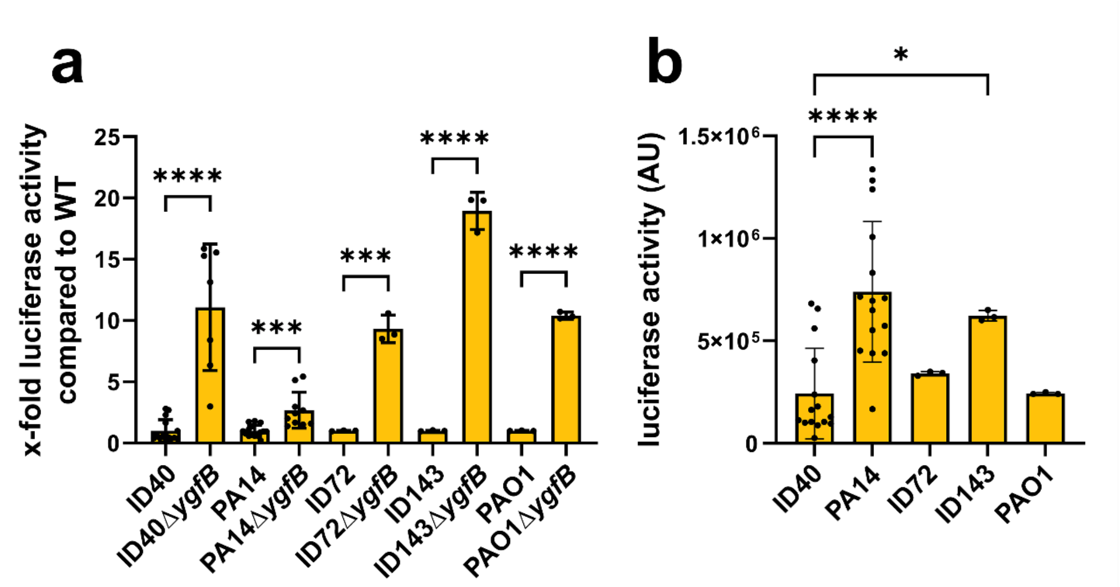

**Fig. S7: Impact of *ygfB* deletion on other *Pa* strains.** (a) and (b) *ampDh3* promoter activity was determined as described in the materials and methods for the indicated strains using the plasmid pBBR harboring the reporter construct *ampDh3*-532-luc comprising the *ampDh3* promoter fragment between position -532 and -1 upstream of the CDS of Nanoluc. Data depict mean and SD. (a) Luciferase activity of *ygfB* deletion strains relative to the "wildtype" is shown using data from *n*=3-15 individual experiments. Asterisks indicate significant differences compared to the "wildtype" strain (*p<0.05, **p<0.1 ***p<0.001, ****p<0.0001; two-tailed Welch's t-test)

(b) Basal promoter activity is shown for the indicated strains using data from *n*=3-15 individual experiments. Asterisks indicate significant differences compared to ID40 (*p<0.05, ****p<0.0001; one-way ANOVA, Dunnett's multiple comparisons comparing to ID40)

**Table S3:** List of strains and plasmids used in this study

| Strain | Relevant characteristics | Source |
|---|---|---|
| ***Pseudomonas aeruginosa*** | | |
| **ID40 strains** | | |
| ID40 | Clinical bloodstream isolate, resistant against cefepime, ceftazidime, imipenem, ciprofloxacin, levofloxacin, piperacillin and piperacillin/tazobactam | [8, 9] |
| ID40Δ*ygfB* | In-frame deletion mutant encoding the first and last 10 amino-acids of *ygfB.* For note: all following deletion mutants are in-frame deletion mutants | [9] |
| ID40Δ*ygfB*::rha-*ygfB* | *ygfB* deletion mutant complemented with pJM220-rha-*ygfB* carrying the *ygfB* coding sequence | [9] |
| ID40Δ*ampDh3* | *ampDh3* deletion mutant | This study |
| D40Δ*ampDh3*::rha-*ampDh3* | *ampDh3* deletion mutant complemented with pJM220-rha-*ampDh3* carrying the *ampDh3* coding sequence | This study |
| ID40Δ*ygfB*Δ*ampDh3* | In frame deletion of both *ygfB* and *ampDh3* | This study |
| ID40Δ*ygfB*Δ*ampDh3*::rha-*ampDh3* | In frame deletion of both *ygfB* and *ampDh3* complemented with pJM220-rha-*ampDh3* carrying the *ampDh3* coding sequence | This study |
| ID40::*ampDh3*-HiBiT | Knock-in mutant carrying a strep-Tag and the 11 amino acid HiBiT-tag at the C-terminal end of AmpDh3 | This study |
| ID40Δ*ygfB*::*ampDh3*-HiBiT | *ygfB* deletion mutant with HiBiT tagged *ampDh3* | This study |
| ID40Δ*ygfB*::rha-*ygfB*:: *ampDh3*-HiBiT | *ygfB* complementant with HiBiT tagged *ampDh3* | This study |
| ID40Δ*alpA* | In-frame deletion mutant of *alpA* encoding the first and last 10 amino-acids of *alpA* | This study |
| D40Δ*alpA*::rha-*alpA* | *alpA* deletion mutant complemented with pJM220-rha-*alpA* carrying the *alpA* coding sequence | This study |
| ID40Δ*ygfB*Δ*alpA* | Double in-frame deletion mutant of *ygfB* and *alpA* | This study |
| ID40Δ*ygfB*Δ*alpA*::rha-*alpA* | Double in-frame deletion mutant of *ygfB* and *alpA* complemented with pJM220-rha-*alpA* | This study |
| ID40::*alpA*-HiBiT | ID40 wildtype with HiBiT tagged *alpA* at the C-terminus | This study |
| ID40Δ*ygfB*::*alpA*-HiBiT | *ygfB* deletion mutant with HiBiT tagged *alpA* at the C-terminus | This study |
| ID40::*alpA*-HiBiT::HA-*alpR* | ID40 wildtype with HiBiT tagged *alpA* at the C-terminus and N-terminal HA-tagged *alpR* | This study |
| ID40Δ*ygfB*::*alpA*-HiBiT::HA-*alpR* | *ygfB* deletion mutant with HiBiT tagged *alpA* at the C-terminus and N-terminal HA-tagged *alpR* | This study |
| ID40Δ*ygfB*::rha-*ygfB* pBBR-532-luc (-532) | *ygfB* complementant with plasmid pBBR-532-luc | This study |
| ID40Δ*ygfB*::rha-*ygfB* pBBR-479-luc (-479) | *ygfB* complementant with plasmid pBBR-479-luc | This study |
| ID40Δ*ygfB*::rha-*ygfB* pBBR-464-luc (-464) | *ygfB* complementant with plasmid pBBR-464-luc | This study |
| ID40Δ*ygfB*::rha-*ygfB* pBBR-430-luc (-430) | *ygfB* complementant with plasmid pBBR-430-luc | This study |
| ID40Δ*ygfB*::rha-*ygfB* pBBR-418-luc (-418) | *ygfB* complementant with plasmid pBBR-418-luc | This study |
| ID40Δ*ygfB*::rha-*ygfB* pBBR-363-luc (363) | *ygfB* complementant with plasmid pBBR-363-luc | This study |
| ID40Δ*ygfB*::rha-*ygfB* pBBR-283-luc (-283) | *ygfB* complementant with plasmid pBBR-283-luc | This study |
| ID40Δ*ygfB*::rha-*ygfB* pBBR-233-luc (-233) | *ygfB* complementant with plasmid pBBR-233-luc | This study |
| ID40Δ*ygfB*::rha-*ygfB* pBBR-180-luc (-180) | *ygfB* complementant with plasmid pBBR-180-luc | This study |
| ID40Δ*ygfB*::rha-*ygfB* pBBR-158-luc (-158) | *ygfB* complementant with plasmid pBBR-158-luc | This study |
| ID40Δ*ygfB*::rha-*ygfB* pBBR-141-luc (-141) | *ygfB* complementant with plasmid pBBR-141-luc | This study |
| ID40Δ*ygfB*::rha-*ygfB* pBBR-122-luc (-122) | *ygfB* complementant with plasmid pBBR-122-luc | This study |
| ID40Δ*ygfB*::rha-*ygfB* pBBR-77-luc (-77) | *ygfB* complementant with plasmid pBBR-77-luc | This study |
| ID40Δ*ygfB*::rha-*ygfB* pBBR-0-luc (0) | *ygfB* complementant with plasmid pBBR-0-luc | This study |

| Strain | Relevant characteristics | Source |
|---|---|---|
| ID40 pBBR-532-luc | ID40 wildtype with plasmid pBBR-532-luc | This study |
| ID40 pBBR-180-luc | ID40 wildtype with plasmid pBBR-180-luc | This study |
| ID40 pBBR-77-luc | ID40 wildtype with plasmid pBBR-77-luc | This study |
| ID40 pBBR-0-luc | ID40 wildtype with plasmid pBBR-0-luc | This study |
| ID40Δ*ygfB* pBBR-532-luc | ID40Δ*ygfB* with plasmid pBBR-532-luc | This study |
| ID40Δ*ygfB* pBBR-180-luc | ID40Δ*ygfB* with plasmid pBBR-180-luc | This study |
| ID40Δ*ygfB* pBBR-77-luc | ID40Δ*ygfB* with plasmid pBBR-77-luc | This study |
| ID40Δ*ygfB* pBBR-0-luc | ID40Δ*ygfB* with plasmid pBBR-0-luc | This study |
| ID40Δ*ygfB*Δ*alpA* pBBR-532-luc | ID40Δ*ygfB* Δ*alpA* with plasmid pBBR-532-luc | This study |
| ID40Δ*ygfB*Δ*alpA* pBBR-180-luc | ID40Δ*ygfB* Δ*alpA* with plasmid pBBR-180-luc | This study |
| ID40Δ*ygfB*Δ*alpA* pBBR-77-luc | ID40Δ*ygfB* Δ*alpA* with plasmid pBBR-77-luc | This study |
| ID40Δ*ygfB* Δ*alpA* pBBR-0-luc | ID40Δ*ygfB* Δ*alpA* with plasmid pBBR-0-luc | This study |
| ID40Δ*ygfB*Δ*alpA*::rha-*alpA* pBBR-532-luc | ID40Δ*ygfB* Δ*alpA* complemented with pJM220-rha-*alpA* with plasmid pBBR-532-luc | This study |
| ID40Δ*ygfB*Δ*alpA*::rha-*alpA* pBBR-0-luc | ID40Δ*ygfB* Δ*alpA* complemented with pJM220-rha-*alpA* with plasmid pBBR-0-luc | This study |
| ID40Δ*alpA* pBBR-532-luc | ID40Δ*alpA* with plasmid pBBR-532-luc | This study |
| ID40Δ*alpA* pBBR-180-luc | ID40Δ*alpA* with plasmid pBBR-180-luc | This study |
| ID40Δ*alpA* pBBR-77-luc | ID40Δ*alpA* with plasmid pBBR-77-luc | This study |
| ID40Δ*alpA* pBBR-0-luc | ID40Δ*alpA* with plasmid pBBR-0-luc | This study |
| ID40Δ*alpA*::rha-*alpA* pBBR-532-luc | ID40Δ*alpA* complemented with pJM220-rha-*alpA* with plasmid pBBR-532-luc | This study |
| ID40Δ*alpA*::rha-*alpA* pBBR-0-luc | ID40Δ*alpA* complemented with pJM220-rha-*alpA* with plasmid pBBR-0-luc | This study |
| ID40::HA-*alpR*::*alpA*-HiBiT pBBR-532-luc | ID40::HA-*alpR*::*alpA*-HiBiT with plasmid pBBR-532-luc | This study |
| ID40::HA-*alpR*::*alpA*-HiBiT pBBR-180-luc | ID40::HA-*alpR*::*alpA*-HiBiT with plasmid pBBR-180-luc | This study |
| ID40Δ*ygfB*::HA-*alpR*::*alpA*-HiBiT pBBR-532-luc | ID40Δ*ygfB*::HA-*alpR*::*alpA*-with plasmid pBBR-532-luc | This study |
| ID40::HA-*alpR*::*alpA*-HiBiT pBBR-180-luc | ID40Δ*ygfB*::HA-*alpR*::*alpA*-with plasmid pBBR-180-luc | This study |
| ID40::rha-*dacB*(PA14) | ID40 strain complemented with pJM220 carrying the *dacB*-coding sequence from PA14 | This study |
| ID40::rha-*dacB*(PA14) PBBR-532-luc | ID40 strain complemented with pJM220 carrying the *dacB*-coding sequence from PA14 | This study |
| ID40Δ*ygfB*::rha-*dacB*(PA14) | Δ*ygfB* complemented with pJM220 carrying the *dacB*-coding sequence from PA14 | This study |
| ID40Δ*ygfB*::rha-*dacB*(PA14):ampDh3-HiBiT | Δ*ygfB* complemented with pJM220 carrying the *dacB*-coding sequence from PA14, *ampDh3* replaced by *ampDH3*-HiBiT | This study |
| ID40Δ*ygfB*::rha-*dacB*(PA14) pBBR-532-luc | Δ*ygfB* complemented with pJM220 carrying the *dacB*-coding sequence from PA14, with plasmid pBBR-532-luc | This study |
| ID40Δ*creBC* | In-frame deletion mutant encoding the first and last 10 amino-acids of the *creBcreC* operon | This study |
| ID40Δ*creBC* pBBR-532-luc | In-frame deletion mutant encoding the first and last 10 amino-acids of the *creBcreC* operon and carrying plasmid pBBR-532-luc | This study |
| ID40 Δ*ygfB* Δ*creBC* | Double deletion mutant of ygfB and of the *creBcreC* operon | This study |
| ID40 Δ*ygfB* Δ*creBC* pBBR-532-luc | Double deletion mutant of ygfB and of the *creBcreC* operon; with plasmid pBBR-532-luc | This study |
| ID40Δ*ampR* | In-frame deletion mutant encoding the first and last 10 amino-acids of *ampR* | This study |
| ID40Δ*ampR* pBBR-532-luc | In-frame deletion mutant encoding the first and last 10 amino-acids of *ampR*; with plasmid pBBR-532-luc | This study |
| **Other *Pa* strains** | | |
| PA14 (DSM No.19882) | Clinical isolate from a human burn patient | DSMZ Braunschweig |
| PA14 pBBR-532-luc | In-frame deletion mutant encoding the first and last 10 amino-acids of *ygfB*; with plasmid pBBR-532-luc | This study |

| Strain | Relevant characteristics | Source |
|---|---|---|
| PA14Δ*ygfB* | In-frame deletion mutant encoding the first and last 10 amino-acids of *ygfB* | This study |
| PA14Δ*ygfB* pBBR-532-luc | In-frame deletion mutant encoding the first and last 10 amino-acids of *ygfB*; with plasmid pBBR-532-luc | This study |
| ID72 | Clinical bloodstream isolate, MDR strain | [8, 10] |
| ID72 pBBR-532-luc | Clinical bloodstream isolate, with plasmid pBBR-532-luc | This study |
| ID72Δ*ygfB* | In-frame deletion mutant encoding the first and last 10 amino-acids of *ygfB* | This study |
| ID72Δ*ygfB* pBBR-532-luc | In-frame deletion mutant encoding the first and last 10 amino-acids of *ygfB;* with plasmid pBBR-532-luc | This study |
| ID143 | Clinical bloodstream isolate, MDR strain | [8] |
| ID143 pBBR-532-luc | Clinical bloodstream isolate, MDR strain; with plasmid pBBR-532-luc | This study |
| ID143Δ*ygfB* pBBR-532-luc | In-frame deletion mutant encoding the first and last 10 amino-acids of *ygfB;* ; with plasmid pBBR-532-luc | This study |
| PAO1 DMSZ No.22644 | Infectious wound | DMSZ Braunschweig |
| PAO1Δ*ygfB* | In-frame deletion mutant encoding the first and last 10 amino-acids of *ygfB;* with plasmid pBBR-532-luc | This study |
| ***Escherichia coli*** | | |
| SM10 λ pir | thi thr leu tonA lacY supE recA::RP4-2-Tc::Mu Km λpir | [11] |
| DH5α | Used for propagation of plasmids during cloning | Thermo Scientific |
| One Shot™ TOP10 Chemically Competent *E. coli* | | Thermo Scientific |
| BL21 DE3 | *fhuA2 [lon] ompT gal (λ DE3) [dcm] ∆hsdS*<br>*λ DE3 = λ sBamHIo ∆EcoRI-B int::(lacI::PlacUV5::T7 gene1) i21 ∆nin5* | New England Biolabs (NEB) |
| **Plasmids** | | |
| pEXG2 | Allelic exchange vector with pBR origin, $Gm^R$, $sacB^+$ | [12] |
| pEXTK | Allelic exchange vector derived from pEXG2, sacB exchanged by thymidine-kinase<br>$Gm^R$, $Tk^+$ | This study |
| pEXG2Δ*ygfB* | pEXG2 derivative for the in-frame deletion of *ygfB*-CDS | [9] |
| pEXG2Δ*creBC* | pEXG2 derivative for the in-frame deletion of *creBC*-CDS | This study |
| pEXG2Δ*ampR* | pEXG2 derivative for the in-frame deletion of *ampR*-CDS | This study |
| pEXG2Δ*alpA* | pEXG2 derivative for the in-frame deletion of *alpA*-CDS | This study |
| pEXG2Δ*ampDh3* | pEXG2 derivative for the in-frame deletion of *ampDh3*-CDS | This study |
| pEXG2::*ampDh3*-HiBiT | pEXG2 derivative for the C-terminal knockin of Strep-HiBiT tag into *ampDh3* | This study made by Genscript |
| pEXG2*::alpA*-HiBiT | pEXG2 derivative for the C-terminal knockin of HiBiT tag into *alpA* | This study,<br>HiBiT tag obtained from Promega |
| pEXTKΔ*ygfB* | pEXTK derivative for the in-frame deletion of *ygfB*-CDS | This study |
| pEXTK::HA-*alpR* | pEXTK derivative for the N-terminal knockin of HA-tag into *alpR* | This study |
| pEXTK::*ampDh3*-HiBiT | pETK derivative for the C-terminal knockin of HiBiT into *ampDh3* | This study |
| pJM220 | Mini-TN7 based vector with transcriptional terminators, rhamnose inducible promoter and MCS, $Gm^R$ | [13] |
| pJM220*ygfB* | pJM220 derivate for the complementation of the *ygfB*-CDS | [9] |
| pJM220*alpA* | pJM220 derivate for the complementation of the *alpA*-CDS | This study |
| pJM220*ampDh3* | pJM220 derivate for the complementation of the *amppDh3*-CDS | This study |
| pJM220*dacB*(PA14) | pJM220 derivate for the complementation of the *dacB*-CDS of PA14 | This study |
| pTNS3 | $Amp^R$, plasmid expressing tnsABCD from P1 and Plac | [14] |
| pFLP2 | $Cb^R$/$Amp^{R;}$ $sacB^+$; Flp recombinase | [15] |
| pGEX-4T3 | Overexpression vector und control of a *tac*-promoter, N-terminal GST tag and a thrombin cleavage site; $Amp^R$ | GE Healthcare |
| pETM-30 | Overexpression vector under the control of a T7-promoter, N-terminal His-GST-tag, N-terminal TEV-cleavage site, $Kan^R$ | EMBL Heidelberg |
| pETM-41 | Overexpression vector under the control of a T7-promoter, N-terminal His-MBP-tag, $Kan^R$ | EMBL Heidelberg |
| pGEX-4T3_ygfB | Derivative of pGEX-4T3 for the overexpression of GST-YgfB | This study |

| Plasmids | Plasmids | Plasmids |
|---|---|---|
| pETM-30_ygfB | Derivative of pETM-30 for the overexpression of His-GST-YgfB | This study |
| pETM-41_alpA | Derivative of pETM-41 for the overexpression of His-MBP-AlpA | This study |
| pNL1.1 | Promoterless basic vector encoding Nanoluc Luciferase | Promega |
| pVT77 | Vector containing lacI-tdk cassette | [16] |
| pME6032 | Modified pVS1-p15A shuttle vector including *lacI*$^{q}$-$P_{tac}$ promoter, $Tc^{r}$ | [17, 18] |
| pME6032-exoU-344 strep-HiBiT | pME6032 derivate encoding ExoU-Strep-HiBiT under control of an exoU promoter replacing *lacI*$^{q}$-$P_{tac}$ promoter | This study |
| pBBR1-MCS-5 | Broad host range vector, Gmr | [19] |
| pBBR-532-luc | pBBR1-MCS-5 derivate containing *ampDh3* promoter fragment bp -532 to -1 upstream of CDS fused to Nanoluc luciferase | This study |
| pBBR-464-luc | pBBR1-MCS-5 derivate containing *ampDh3* promoter fragment bp -479 to -1 upstream of CDS fused to Nanoluc luciferase | This study |
| pBBR-430-luc | pBBR1-MCS-5 derivate containing *ampDh3* promoter fragment bp -430 to -1 upstream of CDS fused to Nanoluc luciferase | This study |
| pBBR-418-luc | pBBR1-MCS-5 derivate containing *ampDh3* promoter fragment bp -418 to -1 upstream of CDS fused to Nanoluc luciferase | This study |
| pBBR-363-luc | pBBR1-MCS-5 derivate containing *ampDh3* promoter fragment bp -363 to -1 upstream of CDS fused to Nanoluc luciferase | This study |
| pBBR-283-luc | pBBR1-MCS-5 derivate containing *ampDh3* promoter fragment bp -283 to -1 upstream of CDS fused to Nanoluc luciferase | This study |
| pBBR-233-luc | pBBR1-MCS-5 derivate containing *ampDh3* promoter fragment bp -233 to -1 upstream of CDS fused to Nanoluc luciferase | This study |
| pBBR-180-luc | pBBR1-MCS-5 derivate containing *ampDh3* promoter fragment bp -180 to -1 upstream of CDS fused to Nanoluc luciferase | This study |
| pBBR-158-luc | pBBR1-MCS-5 derivate containing *ampDh3* promoter fragment bp -158 to -1 upstream of CDS fused to Nanoluc luciferase | This study |
| pBBR-141-luc | pBBR1-MCS-5 derivate containing *ampDh3* promoter fragment bp -141 to -1 upstream of CDS fused to Nanoluc luciferase | This study |
| pBBR-122-luc | pBBR1-MCS-5 derivate containing *ampDh3* promoter fragment bp -122 to -1 upstream of CDS fused to Nanoluc luciferase | This study |
| pBBR-77-luc | pBBR1-MCS-5 derivate containing *ampDh3* promoter fragment bp -77 to -1 upstream of CDS fused to Nanoluc luciferase | This study |
| pBBR-0-luc | pBBR1-MCS-5 derivate containing promoterless Nanoluc luciferase | This study |

#### Table S4. List of the primers used for PCR

| Plasmid | Primer for Gibson Cloning | Sequence 5'–3' |
|---|---|---|
| **pEXG2 derivates**<br>(Gibson assembly of linearized pEXG2 (template pEXG2) with up and down fragment s (template ID40), For validation of mutants the primer GOI_seqF / GOI_seqR and GOI_seqF / GOI_seq_insideR were used | | |
| | gib_uni_pEXG2_f<br>(linearization of vector) | AGGTCGACTCTAGAGGATCC |
| | gib_uni_pEXG2_r<br>(linearization of vector) | TTCCGGCTCGTATAATGTGT |
| **pEXG2ΔcreBC** | 1115creBC_up_F | AGCTAATTCCACACATTATACGAGCCGGAAGATC<br>AGGATCAGCGCGTGTTGAG |
| | 1116creBC_up_R | TCAGCCGCGCGGCAGCCAGAGCAGCGCCTCTTC<br>ATCTTCGAC GATCAGGATATGC |
| | 1117creBC_dn_F | ATGCCGCATATCCTGATCGTCGAAGATGAAGAGG<br>CGCTGCTCTGGCTG |
| | 1118creBC_dn_R | TCGAGCCCGGGGATCCTCTAGAGTCGACCTcAGT<br>CGGCCTTCAGGATGACC |
| | 1119creBC_seq1_F | CAAGGGCTTGCGCAAATTC |
| | 1120creBC_seq2_R | AGGCCGTCGATCATCAAC |
| | 1121creBC_inside_R | GCTTGCAGGCCTCGAAAC |
| **pEXG2ΔampR** | 1213ampR_up_F | AGCTAATTCCACACATTATACGAGCCGGAACACG<br>TCGAGGTGGGTCTG |
| | 1214ampR_up_R | TTATCTCCCCCGCGCCTCAACGGCAGCCATGGC<br>GTTCAGCGGCAAATG |
| | 1215ampR_dn_F | TTGGTTCGACCCCATTTGCCGCTGAACGCCATGG<br>CTGCCGTTGAGGC |
| | 1216ampR_dn_R | TCGAGCCCGGGGATCCTCTAGAGTCGACCTCAG<br>GTGTTCCGCGCCTTC |
| | 1217ampR_seq_F | GTCGTTGGCTGCATGAGAAAC |
| | 1218ampR_seq_R | ATGCTCGAGAGCGAGATCG |
| | 1219ampR_inside_R | CAACCGACCGTGAAGGTTC |
| **pEXG2ΔalpA** | 1572alpA_up_F | AGCTAATTCCACACATTATACGAGCCGGAAAGTA<br>CCCCGAGCATCCAC |
| | 1573alpA_up_R | AATACCATGTTTCAAAGTACCGAGCAGGCGGAG<br>GTGGGGATCGTGGGC |
| | 1574 alpA_dn_F | TTTTTAGCGCTCGCCCACGATCCCCACCTCCGCC<br>TGCTCGGTACTTTG |
| | 1575alpA_dn_R | TCGAGCCCGGGGATCCTCTAGAGTCGACCTTTG<br>AAGCTGCGAATGCGC |
| | 1578AlpA_seq_F | TTGGCTCGGACATGGATG |
| | 1579alpA_seq_R | CAAGCGTCCGTATATGGG |
| | 1580alpA_inside_R | CTGTCGAAGAGCACCCTG |
| **pEXG2ΔcreBC** | 1115creBC_up_F | AGCTAATTCCACACATTATACGAGCCGGAAGATC<br>AGGATCAGCGCGTGTTGAG |
| | 1116creBC_up_R | TCAGCCGCGCGGCAGCCAGAGCAGCGCCTCTTC<br>ATCTTCGAC GATCAGGATATGC |
| | 1117creBC_dn_F | ATGCCGCATATCCTGATCGTCGAAGATGAAGAGG<br>CGCTGCTCTGGCTG |
| | 1118creBC_dn_R | TCGAGCCCGGGGATCCTCTAGAGTCGACCTcAGT<br>CGGCCTTCAGGATGACC |
| | 1119creBC_seq1_F | CAAGGGCTTGCGCAAATTC |
| | 1120creBC_seq2_R | AGGCCGTCGATCATCAAC |
| | 1121creBC_inside_R | GCTTGCAGGCCTCGAAAC |
| **pEXG2ΔampR** | 1213ampR_up_F | AGCTAATTCCACACATTATACGAGCCGGAACACG<br>TCGAGGTGGGTCTG |
| | 1214ampR_up_R | TTATCTCCCCCGCGCCTCAACGGCAGCCATGGC<br>GTTCAGCGGCAAATG |
| | 1215ampR_dn_F | TTGGTTCGACCCCATTTGCCGCTGAACGCCATGG<br>CTGCCGTTGAGGC |
| | 1216ampR_dn_R | TCGAGCCCGGGGATCCTCTAGAGTCGACCTCAG<br>GTGTTCCGCGCCTTC |
| | 1217ampR_seq_F | GTCGTTGGCTGCATGAGAAAC |
| | 1218ampR_seq_R | ATGCTCGAGAGCGAGATCG |
| | 1219ampR_inside_R | CAACCGACCGTGAAGGTTC |

| Plasmid | Primer for Gibson Cloning | Sequence 5'–3' |
|---|---|---|
| **pEXG2ΔalpA** | 1572alpA_up_F | AGCTAATTCCACACATTATACGAGCCGGAAAGTACCCCGAGCATCCAC |
| | 1573alpA_up_R | AATACCATGTTTCAAAGTACCGAGCAGGCGGAGGTGGGGATCGTGGGC |
| | 1574 alpA_dn_F | TTTTTAGCGCTCGCCCACGATCCCCACCTCCGCCTGCTCGGTACTTTG |
| | 1575alpA_dn_R | TCGAGCCCGGGGATCCTCTAGAGTCGACCTTTGAAGCTGCGAATGCGC |
| | 1578AlpA_seq_F | TTGGCTCGGACATGGATG |
| | 1579alpA_seq_R | CAAGCGTCCGTATATGGG |
| | 1580alpA_inside_R | CTGTCGAAGAGCACCCTG |
| **pEXG2ΔampDh3** | 959 ampDH3_up_F | AGCTAATTCCACACATTATACGAGCCGGAACCAGGTGATCCTGTCGATG |
| | 960 ampDH3_up_R | ATGCTGACCATCGACTACAACAGCTATCGCTACGCTCTGAACGAGAAATACCC |
| | 961 ampDH3_dn_F | TCAGGCCGGGTATTTCTCGTTCAGAGCGTAGCGATAGCTGTTGTAGTCGATG |
| | 967ampDh3_dn_R | TCGAGCCCGGGGATCCTCTAGAGTCGACCTTTCATCCTGACCCTCTGCG |
| | 963ampDH3_seq_F | ATTCGGCCATTCGATGAG |
| | 964ampDH3_seq_R | CGGAGGCTTTCCATCATC |
| | 965ampDH3_inside_R | GCCTTCCAGATGCATTTCC |
| **pEXG2::alpA-HiBiT** | 1620alpAHiBiT_up_F | AGCTAATTCCACACATTATACGAGCCGGAAGGACTTGCCGCGCTCCAGATC |
| | 1621alpAHiBiT_up_R | TTCTTGAACAGCCGCCAGCCGCTCACTCCACTCGAACCACCGCTCGAGCCGCGCTCGCCCACGATCCC |
| | 1622alpAHiBiT_dn_F | GAGTGAGCGGCTGGCGGCTGTTCAAGAAGATTAGCTAAAAAAATTTCTGTATGAAAACGGGTTTTCCC |
| | 1623alpAHiBiT_dn_R | GAATTCGAGCTCGAGCCCGGGGATCCTCTAGAGTCGACCTAGGTGCTCGGCACGATAC |
| | 1624alpAHiBiT_seq_F | GGCGGGATTTCGTCCTTG |
| | 1625alpAHiBiT_seq_R | GAGCTTGGCTCGGACATG |
| | 1081HiBiT_inside_R | TCTTCTTGAACAGCCGCC |
| **pEXG2-ampDh3-HiBiT_up** Gibson assembly of linearized pEXG2 with ampDh3up fragment (ID40 as template) and strep-HiBiT fragment | 1072 ampDh3_up_F | AGCTAATTCCACACATTATACGAGCCGGAAATGCTGACCATCGACTACAAC |
| | 1073ampDh3_up_R | CTTCTCAAATTGAGGATGACTCCACGCGCTGGCCGGGTATTTCTCGTT |
| | 1074strep-HiBiT_F | CTCTACGCTCTGAACGAGAAATACCCGGCCAGCGCGTGGAGTCATCC |
| | 1075strep-HiBiT_R | TCGAGCCCGGGGATCCTCTAGAGTCGACCTTTAGCTAATCTTCTTGAACAGCCG |
| | 1073ampDh3_up_R | CTTCTCAAATTGAGGATGACTCCACGCGCTGGCCGGGTATTTCTCGTT |
| **pEXG2-ampDh3-HiBiT** Gibson assembly of linearized pEXG2-ampDh3-HiBiT_up **(pEXG2-ampDh3-HiBiT_up as template)** and downstream fragment | 1076peXG2-HiBiTup_lin_F (Linearization of vector) | TTAGCTAATCTTCTTGAACAGCCGCCAGCCGCTCACTCCACTCGAACCAC |
| | 1077peXG2-HiBiTup_lin_R (Linearization of vector) | AAATCGGCCAGCAGTTGCAGGTTCAGGCCGAGGTCGACTCTAGAGGATCCC |
| | 1078ampDH3dn_F | GGCTGGCGGCTGTTCAAGAAGATTAGCTAAGCCATTAACCGCGAGCG |
| | 1285ampDh3dn_R | TCGAGCCCGGGGATCCTCTAGAGTCGACCTGCTGGTGACCCTGCTCTAC |
| | 1080ampDH3_seq_F | GAAGGAACTCTACGAGGCCG |
| | 1081HiBiT_inside_R | TCTTCTTGAACAGCCGCC |
| | 963ampDh3_seq_R | ATTCGGCCATTCGATGAG |

| Plasmid | Primer for Gibson Cloning | Sequence 5'–3' |
|---|---|---|
| **pEXTK**<br>Gibson assembly with linearized pEXG2 and lacI-tdk fragment from pVT77;1375-1378 sequencing primer | 1371pEXG2_lin_R | AAACGCAAAAGAAAATGCCG |
| | 1372pexAZTlin_F | GCAGGTAAGCTAATTCCACAC |
| | 1373lacI-tdK_F | CTTCCATAATCGGCATTTTCTTTTGCGTTTTTAAT CGTGGCGATGCCTTTC |
| | 1374lacI-tdk_R | TCGTATAATGTGTGGAATTAGCTTACCTGCGACA CCATCGAATGGTGCAAAAC |
| | 1375pEXTKseq1_F | CTTATGTCAATTCGAGAATTACG |
| | 1376pEXTKseq2_R | GAGATTCGTGCGGAACATG |
| | 1377pEXTKseq3_F | ACCTATACGCGAACTGAC |
| | 1378pEXTKseq4_F | GTTAACGGCGGGATATAAC |
| **pEXTK derivates**<br>(Gibson assembly of linearized pEXTK (template pEXTK) with up and down fragment (template ID40). For validation of mutants the primer GOI_seqF / GOI_seq_R and GOI_seq_F / GOI_inside_R were used | | |
| | 1gib_uni_pEXG2_F (linearization of pEXTK vector) | AGGTCGACTCTAGAGGATCC |
| | 1467gib_uni_pEXTK_R (linearization of pEXTK vector) | TTCCGGCTCGTATAATGTGTGGAATTAGCTTACC TGC |
| **pEXTKΔ*ygfB*** (template for ygfB insert derived from pEXG2ΔygfB) | 704pEXG2_ygfB_up_F | AGCTAATTCCACACATTATACGAGCCGGAAgtagca GTCGATCTCGCAG |
| | 707pEXG2_ygfB_dn_R | CTCGCCGAGGCGGCCATGCCGGTCTCGCCGCCT TCACTGCACTGAGGTTTC |
| | 708ygfB_seq_F | CATGACCTTCACTTCGTTG |
| | 709ygfB_seq_R | CTTGTCGAGAATCTGCAC |
| | 710ygfB_inside_R | GGAAGTCATGGAATACCTG |
| **pEXTK-HA-alpR** | 1599HAalpR_up_F | AGCTAATTCCACACATTATACGAGCCGGAAGGAC ATGGATGACTCCCG |
| | 1600HAalpR_up_R | GTAAGCGTAATCTGGAACATCGTATGGGTACATA GGTGGAAAGACTAAGGG |
| | 1601HAalpR_dn_F | TACCCATACGATGTTCCAGATTACGCTTACCCAT ACGATGTTCCAGATTACGCTGAACTCAAAGATCG CATCAAGG |
| | 1602HAalpR_dn_R | TCGAGCCCGGGGATCCTCTAGAGTCGACCTTCA GACCAGCACCGAGTAC |
| | 1603HAlpR_seq_F | GCGAAACGCACGGTCATC |
| | 1604HAalpRseq_R | GCGCTCCAGATCGGAAATG |
| | 1248 HA_ inside_R | TGGAACATCGTATGGGTAAGC |
| **pEXTK-ampDh3-HiBiT**<br>(Gibson assembly using linearized pEXTK and ampDh3-strep-HiBiT fragment (from pEXG2-HiBiT as template) | 1072 ampDh3_up_F | AGCTAATTCCACACATTATACGAGCCGGAAATGC TGACCATCGACTACAAC |
| | 1285ampDh3_dn_R | TCGAGCCCGGGGATCCTCTAGAGTCGACCTGCT GGTGACCCTGCTCTAC |
| | 1080ampDH3_seq_F | GAAGGAACTCTACGAGGCCG |
| | 1081HiBiT_inside_R | TCTTCTTGAACAGCCGCC |
| | 963ampDh3_seq_R | ATTCGGCCATTCGATGAG |
| **pJM220_*dacB* (PA14)**<br>(Gibson assembly of linearized pJM220 with *dacB* coding sequence of PA14 (template PA14) | 773Gib_uni_pJM220_F (linearization of vector) | TACCTCGCGAAGGCCTTGCA |
| | 774Gib_uni_pJM220_R (linearization of vector) | AAGCTTCTCGAGGAATTCCTGC |
| | 1227 pJM220_dacBPA14_F | ATTCAACTAGTGCTCTGCAGGAATTCCTCGAGAA GCTTATGTTCAAGTCGCTGCGTAC |
| | 1228pJM220_dacB PA14_R | CTGGTTGGCCTGCAAGGCCTTCGCGAGGTATTAT TTCCGCGCGTGCAG |
| **pJM220_*alpA***<br>(Gibson assembly of linearized pJM220 with *dacB* coding sequence of PA14 (template PA14) | 773Gib_uni_pJM220_F (linearization of vector) | TACCTCGCGAAGGCCTTGCA |
| | 774Gib_uni_pJM220_R (linearization of vector) | AAGCTTCTCGAGGAATTCCTGC |
| | 2074 pJM220_alpA_F | AGTGCTCTGCAGGAATTCCTCGAGAAGCTTATGT TTCAAAGTACCGAG |
| | 2075pJM220_alpA_R | CTGGTTGGCCTGCAAGGCCTTCGCGAGGTATTA GCGCTCGCCCACGATC |

| **Plasmid** | **Primer for Gibson Cloning** | **Sequence 5'–3'** |
|---|---|---|
| **pJM220_*ampDh3*** (Gibson assembly of linearized pJM220 with *dacB* coding sequence of PA14 (template PA14) | 773Gib_uni_pJM220_F (linearization of vector) | TACCTCGCGAAGGCCTTGCA |
| | 774Gib_uni_pJM220_R (linearization of vector) | AAGCTTCTCGAGGAATTCCTGC |
| | 2072pJM220_ampDh3_F | AGTGCTCTGCAGGAATTCCTCGAGAAGCTTATGCTGACCATCGACTACAAC |
| | 2073pJM220_ampDh3_R | CTGGTTGGCCTGCAAGGCCTTCGCGAGGTATCAGGCCGGGTATTTCTCG |
| **pGEX4T3_ygfB** (Gibson assembly of linearized pGEX4T3 with coding sequence of *ygfB* (template ID40); 847 and 848 sequencing primer) | 845Gib_pGEX4T3_R | GGAATTCGGGGATCCACGCG |
| | 846Gib_pGEX4T3_F | CGGGTCGACTCGAGCGGC |
| | 843Gib_pGEX4T3_ygfB_F | GATCTGGTTCCGCGTGGATCCCCGAATTCCATGCCGGTCTCGCCGGCC |
| | 844Gib_pGEX4T3_ygfB_R | ATCGTCAGTCAGTCACGATGCGGCCGCTCGAGTCGACCCGTCAGTGCAGTGAAGGCTTGGGAGCGG |
| | 847pGEX4T3_seq_F | ATGTTGTATGACGCTCTTGA |
| | 848pGEX4T3_seq_R | CAAGAATTATACACTCCGCT |
| **pETM-41_alpA** (Gibson assembly of linearized pETM-41 with coding sequence of *alpA* (template ID40)) | 1769 pETM41_lin_F | TAATGAGGTACCGGATCCGAATTCGAGC |
| | 1770 pETM41_lin_R | GCCCATGGCGCCCTGAAAATAAAG |
| | 1771_pETM41_MBP-AlpA-fw | GAGAATCTTTATTTTCAGGGCGCCATGGGCTTTCAAAGTACCGAGCAGGCG |
| | 1772_pETM41_MBP-AlpA_rw | GAGCTCGAATTCGGATCCGGTACCTCATTATTAGCGCTCGCCCACGAT |
| **pETM-30_ygfB** (Gibson assembly of linearized pETM-30 with coding sequence of *ygfB* (template ID40)) | 1824_pETM-30_linF | TAAGGTACCGGATCCGAATTCG |
| | 1825_pETM-30_linR | GCCATGGCGCCCTGAAAA |
| | 1826_pETM-30_ygfB_insert_F | GAGAATCTTTATTTTCAGGGCGCCATGGCGATGTCCACTCAGAATTCCGCC |
| | 1827_pETM-30_ygfB_insertR | ACGGAGCTCGAATTCGGATCCGGTACCTTAGTGCAGTGAAGGCTTGGG |
| **pME6032-exoU_D344A-Strep-HiBiT** Gibson assembly of linearized pME6032 and upstream and downstream fragment. The upstream fragment was generated by PCR with 167 and 168 (template PA14). Downstream fragment was stepwise generated by amplifying first PA14 template with 169 and 163. The PCR product was used for reamplification with 169 and 164. This PCR product was again amplified with 169 and 170 to obtain the downstream fragment. | 165_Gib_uni_pME6032F | GGATCCGATATCGCCGTGGC |
| | 166_Gib_uni-pME6032R | GAATTCGAGCTCCCGGGTAC |
| | 167 exoU_up_F | CGGTGAGAATGGCAAAAGCCGCCACGGCGATATCGGATCCCCTACTAACTAGGCAGCGGTAC |
| | 168exoU_up_R | cgttaatcatcaccccgccaGCCTGGAATTCTGTCCACTC |
| | 169exoUstrep_dn_F | GAGTGGACAGAATTCCAGGCTGGCGGGGTGATGATTAACG |
| | 163exoUstrep_dn_R | TTTTTCGAACTGCGGGTGGCTCCACGCGCTCTTCTCAAATTGAGGATGACTCCACGCGCTTGTGAACTCCTTATTCCGCCAAG |
| | 164exoUstrep-HiBiT_R | TTAGCTAATCTTCTTGAACAGCCGCCAGCCGCTCACTCCACTCGAACCACCGCTCGAGCCTTTTTCGAACTGCGGGTGGC |
| | 170exoU_dn_R | ATCTATCGATGCATGCCATGGTACCCGGGAGCTCGAATTCTTAGCTAATCTTCTTGAACAGCCG |
| **pBBR-532-luc** Gibson assembly of linearized pBBR with ampDh3 promoter fragment -1 to -532 prior to CDS (template ID40) and nanoluc (template pNL1.1) | 946 pBBR1_lin_R | GCGTTAATATTTTGTTAAAATTCGCG |
| | 947 pBBR1_lin_F | GCCTGGGGTGCCTAATGAG |
| | 948 nanoluc pBBR-F | AACGCGAATTTTAACAAAATATTAACGCTTACGCCAGAATGCGTTCG |
| | 949nanoluc_pbbr R | CTTTCCATCATCCGAAAAAAGGTGAAAACCATGGTCTTCACACTCGAAGATTTC |
| | 950ampDH3_532_ F | CCCAACGAAATCTTCGAGTGTGAAGACCATGGTTTTCACCTTTTTTCGGATGATG |
| | 951 ampDH3_532_R | GAGTTAGCTCACTCATTAGGCACCCCAGGCTAGCCGCTCTGTCGAGGG |
| **pBBR-532-luc derivates** Gibson assembly of linearized pBBR with various ampDh3 promoter-nanoluc fragments using pBBR-532-luc as template and for amplification 948 as forward primer and different reverse primers as depicted | | |
| pBBR_479-luc | 1035ampDH3_479_R | GAGTTAGCTCACTCATTAGGCACCCCAGGCCAAAGTAGAGCGGCCAACG |
| pBBR_464-luc | 1041ampDh3_464_R | GAGTTAGCTCACTCATTAGGCACCCCAGGCAACGGACGAACGGTGTTG |

| **Plasmid** | **Primer for Gibson Cloning** | **Sequence 5‘–3‘** |
|---|---|---|
| pBBR_430-luc | 1034ampDh3_430_R | GAGTTAGCTCACTCATTAGGCACCCCAGGCCGC GGTAGTTTTTTCCCATGATC |
| pBBR_418-luc | 968ampDh3_418_R | GAGTTAGCTCACTCATTAGGCACCCCAGGCTTCC CATGATCACGTTTCCCG |
| pBBR_363-luc | 1042ampDh3_363_R | GTGAGTTAGCTCACTCATTAGGCACCCCAGGCTT CGTCGTTTCCGCCCG |
| pBBR_283-luc | 1043ampDh3_283_R | GAGTTAGCTCACTCATTAGGCACCCCAGGCTCC GGCCAACTTAAAGTTCGG |
| pBBR_233-luc | 969ampDh3_233_R | GAGTTAGCTCACTCATTAGGCACCCCAGGCTTCA GTGGCCTCGGTGAG |
| pBBR_180-luc | 1044ampDh3-180_R | GAGTTAGCTCACTCATTAGGCACCCCAGGCGTGT ATCCCTTTTGCCCGGC |
| pBBR_141-luc | 1045ampDh3-141_R | GAGTTAGCTCACTCATTAGGCACCCCAGGCTGC CCGGTCTTCTTCCTTC |
| pBBR_122-luc | 1033ampDh3-122_R | GAGTTAGCTCACTCATTAGGCACCCCAGGCCGG TTTAGCCGGGGCCC |
| pBBR_77-luc | 1045ampDh3-141_R | GAGTTAGCTCACTCATTAGGCACCCCAGGCATCG CTGCATCGACGTCC |
| pBBR_0-luc | 1032ampDh3-0_R | GAGTTAGCTCACTCATTAGGCACCCCAGGCATG GTCTTCACACTCGAAGATTTC |
| **Primer for qRT PCR** | **Name** | **Sequence 5‘–3‘** |
| | alpB_F | CGTAATAGCCGCCGACCC |
| | alpB_R | CGCCATCTTCTTTGTCGTGT |
| | alpC_F | TGCTCGGCACGATACAACC |
| | alpC_R | CGATCTGCATGCGCCTGAT |
| | alpD_F | TAGTAAGCCGTTCCACCCGT |
| | alpD_R | TCGACCTTGTTCTCGCTCAC |
| | alpE_F | GCATGGAAAAGACGGACGAC |
| | alpE_R | TTGAGCCTGACCATAACCGC |
| | ampC_F | GAAGGCTACGGGGTGAAGAC |
| | ampC_R | CGACCTTGTAGTAACCGCGA |
| | ampD_F | GGAGACGAACTGGGTGATGG |
| | ampD_R | GGTAAGGTCCAGGCGTTCTT |
| | ampDh2_F | CCGAGGTGTAGTCGGTGTTC |
| | ampDh2_R | GCGTTCGTTCGTTCTACTGG |
| | ampDH3_F | TACCGGCCTCGTAGAGTTCC |
| | ampDH3_r | CACTCAAGCAACTGGCGAAG |
| | rpoS_F | TGCCGATCCATGTGGTCAAG |
| | rpoS_R | GTTGGCGATTTCTTCGGGTG |
| | TUEID40_01954_F | TGTTGCGGATGCTGACGAA |
| | TUEID40_01954_R | GAATGGCGACAGCAACCCTT |
| | ygfB_F | TCCATATAGTCCGTCTCGCC |
| | ygfB_R | GGGTTTCCTCGCCGGTTT |

#### Table S5: UniProt accession numbers of the protein sequences aligned in Fig. S1.

| **Species** | **Name** | **Accession** |
|---|---|---|
| *Acinetobacter baumannii* (Ab) | YgfB/YecA family protein | V5VAK1 |
| *Escherichia coli* (strain K12) (Ec K12) | UPF0149 protein YgfB | P0A8C4 |
| *Haemophilus influenzae* (strain ATCC 51907 / DSM 11121 / KW20 / Rd) (Hi ATCC 51907) | UPF0149 protein HI_0817 | P44882 |
| *Klebsiella pneumoniae subsp. pneumoniae* (strain ATCC 700721 / MGH 78578) (Kp ATCC 700721) | UPF0149 protein KPN78578_32810 | A6TDS1 |
| *Legionella pneumophila subsp. pneumophila* (strain Philadelphia 1 / ATCC 33152 / DSM 7513) (Lp ATCC 33152) | lpg0076 protein | Q5ZZD5 |
| *Pasteurella multocida* (strain Pm70) (Pm Pm70) | UPF0149 protein PM1723 | Q9CKA2 |
| *Pseudomonas aeruginosa* (strain UCBPP-PA14) (Pa PA14/ID40) | UPF0149 protein PA14_69010 | Q02ED9 |
| *Pseudomonas fluorescens* (strain ATCC BAA-477 / NRRL B-23932 / Pf-5) (Pf ATCC BAA-447) | UPF0149 protein PFL_5969 | Q4K406 |
| *Pseudomonas putida* (strain ATCC 47054 / DSM 6125 / NCIMB 11950 / KT2440) (Pp ATCC 47054) | UPF0149 protein PP_5201 | Q88CI0 |
| *Salmonella typhimurium* (strain LT2 / SGSC1412 / ATCC 700720) (St ATCC 700720) | UPF0149 protein YgfB | Q8ZM71 |
| *Shigella flexneri* (Sf) | UPF0149 protein YgfB | P0A8C7 |
| *Vibrio cholerae* serotype O1 (strain ATCC 39315 / El Tor Inaba N16961) (Vc ATCC 39915) | UPF0149 protein VC_2476 | Q9KP97 |
| *Xanthomonas campestris pv. campestris* (strain ATCC 33913 / DSM 3586 / NCPPB 528 / LMG 568 / P 25) (Xc ATCC 33913) | UPF0149 protein XCC3260 | Q8P5S6 |
| *Yersinia enterocolitica* serotype O:8 / biotype 1B (strain NCTC 13174 / 8081) (Ye 8081) | UPF0149 protein YE3397 | A1JPP0 |
| *Yersinia pestis* (Yp) | UPF0149 protein PO0911/y3298/YP_3608 | Q8ZHI2 |
| *Yersinia pseudotuberculosis* serotype O:3 (strain YPIII) (Yps) | UPF0149 protein YPK_0862 | B1JNS1 |
